# Brain structure and activity predicting cognitive maturation in adolescence

**DOI:** 10.1101/2024.08.23.608315

**Authors:** Junda Zhu, Clément M. Garin, Xue-Lian Qi, Anna Machado, Zhengyang Wang, Suliann Ben Hamed, Terrence R. Stanford, Emilio Salinas, Christopher T. Whitlow, Adam W. Anderson, Xin Maizie Zhou, Finnegan J. Calabro, Beatriz Luna, Christos Constantinidis

## Abstract

Cognitive abilities of primates, including humans, continue to improve through adolescence ^1,2^. While a range of changes in brain structure and connectivity have been documented ^3,4^, how they affect neuronal activity that ultimately determines performance of cognitive functions remains unknown. Here, we conducted a multilevel longitudinal study of monkey adolescent neurocognitive development. The developmental trajectory of neural activity in the prefrontal cortex accounted remarkably well for working memory improvements. While complex aspects of activity changed progressively during adolescence, such as the rotation of stimulus representation in multidimensional neuronal space, which has been implicated in cognitive flexibility, even simpler attributes, such as the baseline firing rate in the period preceding a stimulus appearance had predictive power over behavior. Unexpectedly, decreases in brain volume and thickness, which are widely thought to underlie cognitive changes in humans ^5^ did not predict well the trajectory of neural activity or cognitive performance changes. Whole brain cortical volume in particular, exhibited an increase and reached a local maximum in late adolescence, at a time of rapid behavioral improvement. Maturation of long-distance white matter tracts linking the frontal lobe with areas of the association cortex and subcortical regions best predicted changes in neuronal activity and behavior. Our results provide evidence that optimization of neural activity depending on widely distributed circuitry effects cognitive development in adolescence.

## Introduction

Maturation of executive functions including working memory is a hallmark of human cognitive development ^1,6^. Delayed response tasks reveal an improvement in performance throughout childhood and adolescence in both precision and reaction time^7–9^ . Although such tasks are simple conceptually, enhancement in their performance proceeds in precise tandem with improvement across a range of other cognitive domains, including response inhibition, task switching, and planning suggesting a domain-general process of cognitive maturation ^10^. Age-related cognitive improvement coincides with brain structural changes, including inverted U-shaped trajectories of total brain volume, driven by decreasing gray matter volumes in adolescence, while white matter volumes continue to increase into adulthood ^3^ . An overall pattern of decreased thickness in prefrontal cortex is evident into adulthood ^5,11^ and is thought to represent restructuring processes such as pruning of infrequently used synapses ^12–14^ . Changes in prefrontal cortex and its connections with other regions continue to progress late in adolescence ^15,16^. Myelination in the underlying white matter providing connectivity between frontal and other regions correlates with development of working memory and other cognitive functions^17,18^. Abnormalities in such processes have been associated with psychopathology of mental illnesses that emerge at the end of adolescence, such as schizophrenia ^19,20^.

The changes in brain activity that underlie cognitive development through adolescence have been addressed primarily with fMRI studies, which reveal distinct differences in prefrontal activity patterns between childhood and adulthood in humans during working memory tasks ^21,22^. More direct evidence about changes in activity of single neurons and populations has been obtained from non-human primate models of adolescence ^23,24^. Rhesus monkeys (*Macaca mulatta*) enter puberty at approximately 3.5 years of age and reach full sexual maturity at 5, aging at a rate of approximately 3 times faster than humans ^25,26^. Anatomical studies suggest a protracted period of prefrontal cortical development that parallels that of humans ^27–29^. Cortical thickness, surface area, and white matter myelination also appear to follow similar trajectories to those of humans, at least in childhood ^30^ ^31^. Biochemical and anatomical changes have also been characterized in the monkey prefrontal cortex from pre-puberty to adulthood, including changes of interneuron morphology and connections ^32,33^.

However, a direct link between cognitive performance in terms of underlying changes in neural activity and concomitant structural brain changes has been lacking. We were therefore motivated to track behavior, neuronal activity, and anatomical imaging measures at multiple developmental stages with key developmental milestones in a cohort of developing monkeys. By determining the relationship of such measurements, we identify the brain mechanisms that can account for the cognitive improvement seen during this critical period of life.

## Results

We tracked developmental measures longitudinally in a cohort of eight monkeys (Group A, six males, two females) on a quarterly basis, in tandem with neurophysiological recordings and MR imaging, from an age of 3.0±0.1 to 7.1±0.1 years (corresponding to human ages of ∼9-21 years). Another six older monkeys matched for training time in the task at different time points were used for control comparisons (groups B and C). At the beginning of the study, morphometric measures were all consistent with individuals in a growth trajectory, however, the emergence of developmental markers and secondary sexual characteristics, such as the eruption of canines, varied considerably between individuals (Extended Data Fig. 1). This reflects the variability in pubertal timing, which defines the period of adolescence. We therefore sought to align individual growth trajectories on a biological developmental marker rather than chronological age and relied on the closure of the epiphysial growth plate, a well-established indicator of skeletal maturation in humans ^34,35^. Thus, we defined a “mid-adolescence” age for each monkey as the time of the tibial epiphyseal closure (see methods). We also examined the developmental trajectory for each morphometric measure on chronological age and mid-adolescence age independently using non-linear regression models (general additive mixed models - GAMM). We compared models aligned on chronological age and mid-adolescence age for each of 7 morphometric measures using the Akaike Information Criterion (AIC). All but one morphometric measure had lower AIC when using mid-adolescence age, compared with using chronological age (table S1). For example, plotting canine length as a function of age across individuals generated a deceptively monotonic curve throughout adolescence and into adulthood (Extended Data Fig. 1b). Instead, aligning on mid-adolescence age revealed that lengthening of canines was completed 21.7 months after the mid-adolescence age on average (Extended Data Fig. 1c). Mid-adolescence age thus provided a more accurate representation of each animal’s developmental stage compared to their chronological age allowing us to effectively account for the individual variability and more directly examine the effects of maturation (Extended Data Fig. 1b-c). The mean mid-adolescence age across individuals was 57.9±3.6 months (corresponding to a human age of ∼14.5 years). All morphometric measures exhibited significant development-related differences and followed a similar, non-linear developmental trajectory, with rapid changes before and around mid-adolescence age, before plateauing when adulthood was reached (Extended Data Fig. 1c-h).

### Working memory performance and developmental measures follow a common trajectory

We evaluated working memory performance with variants of the oculomotor delayed response task (Fig. 1a-c), which has been extensively used in human studies ^7,10^. Monkeys were required to observe a visual cue that could appear at one of eight locations and, after a delay period, to make an eye movement to the location of the remembered visual stimulus. A distractor stimulus appeared in a variant of the task (ODR + distractor). Behavioral performance was collected at time points spaced ∼4 months apart from 3.4 to 6.2 years old. Animals were able to perform the task from their earliest time point and achieved an average of 79% correct responses (excluding trials aborted before the end of the delay period – Extended Data Fig. 2), in agreement with previous studies in monkeys ^23^ and human children ^8^. Human studies have revealed that the precision of working memory improves in adolescence^7,8,10^. We similarly sought to capture changes in working memory functions beyond simply task performance.

**Figure 1.**
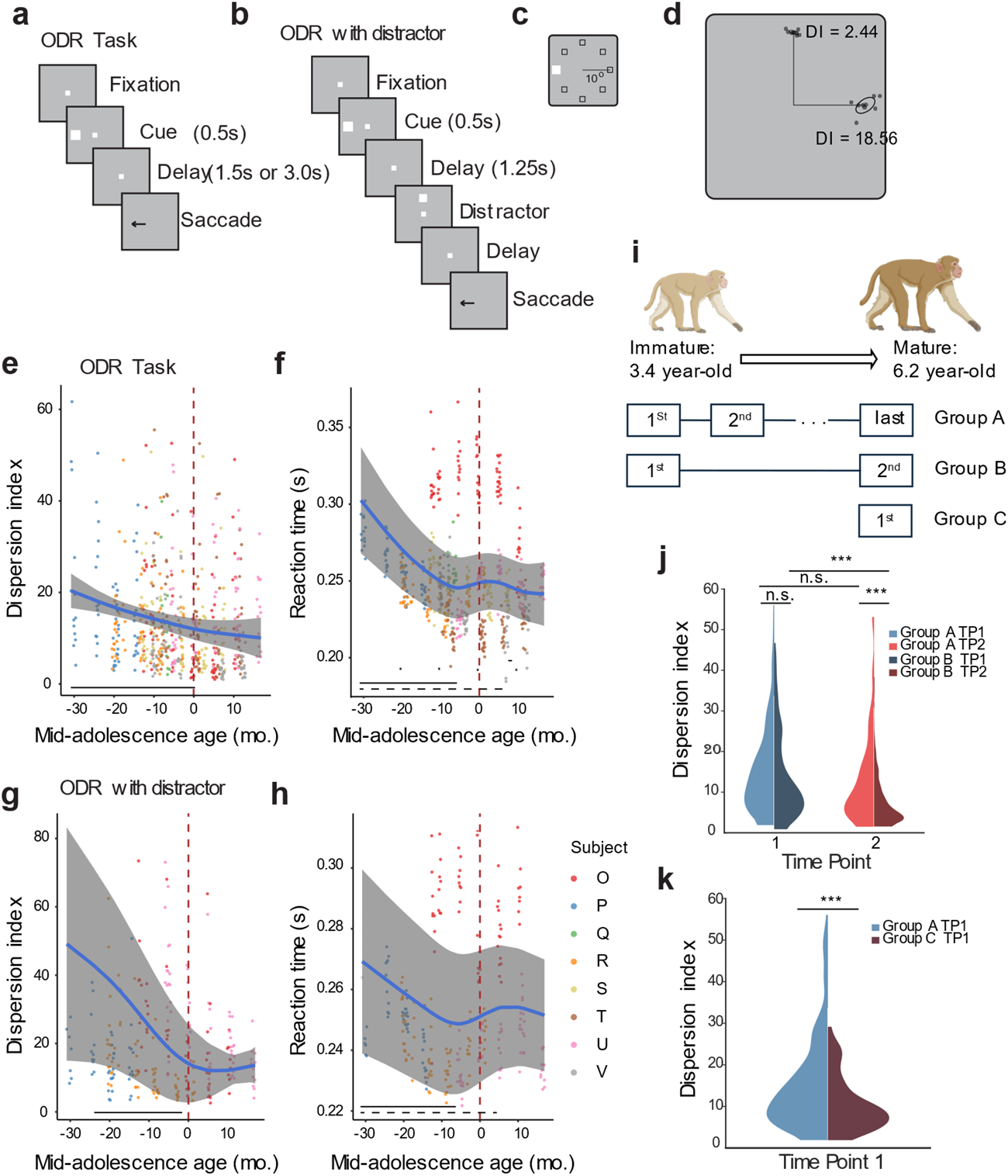
Saccade precision and latency improve during adolescence. (a) Sequence of events in the ODR task. The monkey is required to maintain fixation while a cue stimulus is presented and after a delay period, when the fixation point turns off, saccade to the remembered location of the cue. (b) Sequence of events in the ODR with distractor task. After the delay period, a distractor stimulus appears, which needs to be ignored. The monkey is still required to saccade to the remembered location of the cue. (c) Possible locations of the stimulus presentation on the screen. (d) Schematic illustration of variability of two groups of saccades. Gray dots represent endpoints of individual saccades for two stimulus locations. Dispersion Index (DI), defined as the area within one standard deviation from the average landing position of each target is shown. (e) DI in the ODR task, during neural recording sessions. Each dot is one session; data from different monkeys are shown in different colors. Blue line shows the GAMM fitted trajectory. Gray shaded regions denote the 95% confidence intervals (CIs). Dashed vertical line denotes the mid-adolescence age 0. The horizontal dashed bar denotes significant developmental effect intervals. The horizontal solid bar denotes significant developmental effect intervals. (f) As in e, for reaction time of saccade in the ODR task. (g) As in e, for DI in the ODR with distractor task. (h) As in f, for reaction time in the ODR with distractor task. (i) Schematic diagram of the three cohorts of monkeys (Groups A-C) used to evaluate behavioral improvement. (j) DI in the ODR task of Group A and B at first (TP1) and second time points (TP2). The violin plot shows the distribution of DI values for both groups at two distinct time points, with the width of the plot indicating the density of data points. (k) DI in the ODR task of Group A and C at the first time point. (*** p < 0.0001).

We therefore used the dispersion index (DI - Fig. 1d) to quantify precision of saccadic endpoints (see Methods), and relied on GAMMs to examine development-related effects. Similar to prior findings in humans using the same task ^36^, working memory precision improved with maturation (GAM, F(1, 568) = 3.31, p = 0.0002; Fig. 1e). Additionally, we examined response latency by calculating the reaction time (RT) of each trial. Importantly, reaction time became faster with maturation indicating that improved precision was not achieved as a result of speed-accuracy tradeoff (GAM, F(1, 568) = 7.76, p = 1.41e-06; Fig. 1f). Working memory precision and latency also improved with age for the variant of the working memory task involving a distractor (DI GAM, F(1, 238) = 2.09, p = 0.0001; RT GAM, F(1, 238) = 5.96, p = 3.47e-05, Fig. 1g-h). In both tasks, significant developmental improvements were found to occur in early adolescence (-31 to 0 months relative to the mid-adolescence age). Reaction time was found to have significant growth rates throughout adolescence (-31 to -7 months relative to the mid-adolescence age), with greater changes earlier. In general, rapid and large developmental improvements were found to occur before and around mid-adolescence age of the monkeys for both saccade precision and latency, followed by a slowdown of change and plateau in late adolescence and early adulthood.

The monkeys tested later in development had more cumulative exposure to the task than in early age, as an inevitable consequence of our longitudinal experimental design. To evaluate the effect of exposure to the task, we compared the behavioral performance of this group of animals (Group A) with 2 other groups of animals’ performance following similar training and developmental procedures (Fig. 1i). We reanalyzed a group with 4 animals (Group B, all males) that was introduced to the same task at a similar starting age (median 4.3 years) and were trained under the same protocol ^23^. After completing their first time point (young stage), their second time point for behavioral testing began 1.6–2.1years later. We compared the saccade precision and its changes in the two groups of animals during their respective 1^st^ and 2^nd^ testing time points. Group B animals had slightly lower DI than the Group A animals during first testing time point that did not reach statistical significance, considering they are slightly older than Group A monkeys (Mean DI of Group A = 15.36, Mean DI of Group B = 13.66, p = 0.21, two-sample t test; Fig. 1j). When comparing the DI in the second time point, we saw a significant difference in the saccade precisions between the groups even though the two groups had comparable exposure in the task (Mean DI of Group A = 13.15, Mean DI of Group B = 7.59, p = 2.83e-04, two-sample t test; Fig. 1j). The result indicates that the difference of age during the two testing time points in two groups accounted for significant difference in saccade precisions in the second time points. We then proceeded to train and test a third group (Group C) of two animals that were first trained to perform the task as adults (6.5 y.o.), with the same training procedure. Adult animals achieved a significantly lower DI in the ODR task despite having the same training exposure as the young animals during their first testing time point (Mean DI of Group A = 15.36, Mean DI of Group C = 10.54, p = 9.35e-05, two-sample t test; Fig. 1k). These results indicated that development confers an improvement in cognitive function and that this cannot be accounted for solely by cumulative task experience.

### Reliability of firing metrics of PFC neurons improves with adolescent development

Neurophysiological recordings were made in areas 46 and 8a of the dorsolateral prefrontal cortex (Fig. 2a), from each animal. To make an unbiased comparison of PFC neural activity, we recorded all neurons we encountered. For each neuron, we determined an age relative to the mid-adolescence age of the animal at the day the neuron was recorded.

**Figure 2.**
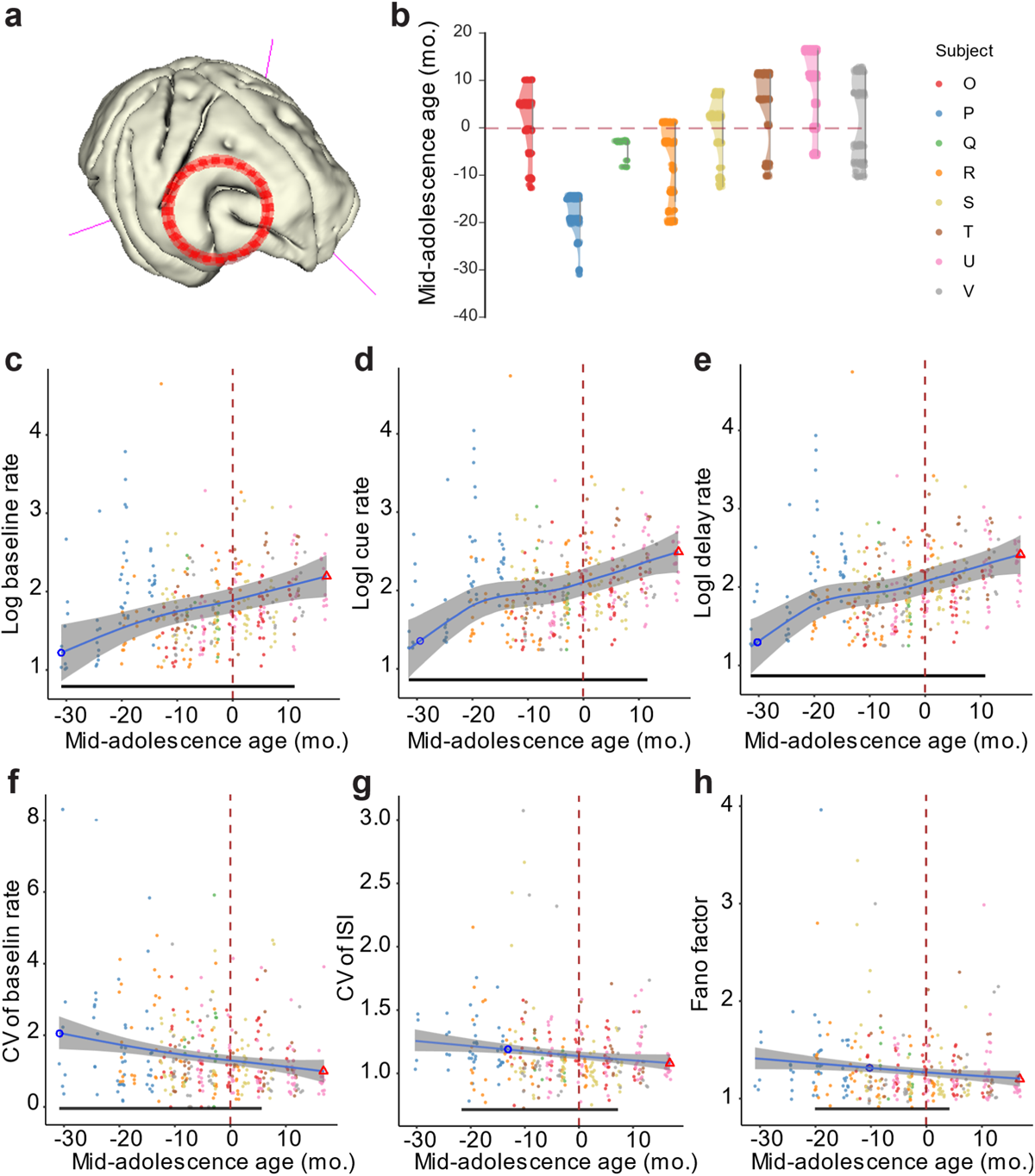
Neurophysiological recordings and neuron firing metrics. (a) Reconstructed MRI T1 image of one monkey’s brain with the placement of recording chamber indicated. (b) Distribution of neurons recorded across time points. Each point represents a neuron at its recorded time to mid-adolescence age shown in the y-axis. Each vertical distribution represents one subject. (c-e) Lognormal average baseline, cue-, and delay-period firing rate of neurons in each session recorded during ODR task. Each dot is one session. Blue line shows the GAMM fitted trajectory. Gray shaded regions denote the 95% confidence intervals (CIs). Blue circle denotes the time of peak development velocity. The red triangle denotes the time of maximum or minimum value. Black bar indicates regions of significant development-related change. (f – h) Coefficient of variation (CV) of firing rate, CV of Inter-spike intervals and Fano factor of neurons in each session recorded during ODR task. The dashed vertical line denotes the mid-adolescence age 0. The horizontal bar denotes significant developmental effect intervals.

In total, our database contained 2131 neurons from 387 sessions across 8 animals (table S2), covering an extended range of adolescence (Fig. 2b). We first determined what aspects of PFC firing rate change over the course of development that could account for the changes we observed in working memory task performance. PFC activity during the baseline (pre-cue) fixation, visual response (cue) and delay epoch changed along with maturation and each followed a trajectory similar to the common trajectory body growth and cognitive function shared, characterized by a significant increase early on, continuing to increase after the mid-adolescence age (Fixation period: F(1, 378) = 5.9, p = 3.4e-06, GAM; significant developmental improvements -31 to 11 months relative to mid-adolescence age; Fig. 2c; Cue period: F(1, 378) = 4.4, p = 0.00023, GAM; significant developmental improvements -31 to 11 months relative to mid-adolescence age; Fig. 2d; Delay period: F(1, 378) = 4.7, p = 0.00020, GAM; significant developmental improvements -31 to 11 months relative to mid-adolescence age; Fig. 2e). We computed the correlation between the GAMM fitted trajectories of behavioral measures and baseline activity of PFC neurons, which revealed a remarkable correlation of neural activity with saccadic precision (DI vs. baseline activity: r = -0.997, permutation test, p<0.0001, Extended Data Fig. 17a) and also strong correlation with reaction time (RT vs. baseline activity: r = -0.930, permutation test, p<0.0001, Extended Data Fig. 17b).

Since the improvement in behavior was to a large extent a decrease in variability, we sought to test whether measures of neural firing variability also declined. We therefore calculated the coefficient of variation (CV) of inter-spike interval and firing rate across trials during pre-cue fixation, which represents the baseline activity, to investigate the intrinsic firing properties of neurons at different developmental stages. Both CV of ISI and baseline firing rate significantly decreased before stabilizing at the adult level (CV of firing rate: F(1, 378) = 3.68, p = 0.00014, GAM; significant developmental improvements -31 to 6 months relative to mid-adolescence age, Fig. 2f; CV of ISI: F(1, 378) = 2.16, p = 0.002, GAM; significant developmental improvements -21 to 7 months relative to mid-adolescence age, Fig. 2g). Similarly, we tested the Fano factor of spike counts. Overall, Fano factor values were significantly lower in the adult than in the young prefrontal cortex (Fig. 2h), and the changes along the maturation resembled the trajectory observed in firing rates (F(1, 326) = 1.8, p = 0.0049, GAM; significant developmental improvements -19 to 5 months relative to mid-adolescence age).

Other aspects of neuronal firing remained relatively stable in adolescence. These included the width of tuning curves for the eight cue locations measured during the cue and delay period. The tuning width did not show significant changes during adolescence in either the cue (p = 0.109, GAM) or delay (p = 0.271, GAM) period. The result did not change after eliminating neurons with low firing rate (baseline ≤0.5 Hz) and poor Gaussian fitting (R^2^ < 0.5) (cue: p = 0.273, GAM in Extended Data Fig. 3a; delay: p = 0.091, GAM in Extended Data Fig. 3b). We also quantified the amount of information carried in single neurons about the location of the cue. Here, we used the percentage of explained variance (ωPEV) statistic to measure the extent to which the variability in neural firing rate in different trials was explained by cue location during stimulus presentation. Overall, PFC neurons were selective for cue locations during the cue period (session average ω^2^ = 0.034, permutation test, p = 0.003) and delay period (session average ω^2^ = 0.033, p = 0.012). We then examined the developmental effect on single neuron stimuli selectivity. ωPEV during visual cue or memory delay did not change significantly during adolescence (p = 0.995, Extended Data Fig. 3c). We assessed whether this averaged measure is driven by subpopulations of neurons that are task responsive. Task responsive neurons had higher ωPEV than the whole population, but there was not significant development effect on ωPEV during visual cue (p = 0.065, GAM) or memory delay (p = 0.372, GAM). We additionally tested whether the results for the whole population were affected by neurons with low firing rate during certain task epochs. The result did not change even after eliminating neurons with low mean firing rate in each epoch (≤0.5 Hz). The intrinsic timescale is another fundamental property that reflects how quickly a neuron responds to changes in input or internal states. The intrinsic timescales remained stable during the adolescent period (p = 0.227, GAM; Extended Data Fig. 3d).

### Dimensionality of PFC responses increases in adolescence

Measures of neural activity examined so far relied on mean values computed over entire task epochs. However, dynamics of neural activity vary at a finer time scale, and we wished to examine how the representation of task conditions may vary in neural responses over time. We employed a method to evaluate the effective temporal coding dimensionality of each neuron using principal component analysis (PCA; see Methods). We then determined the effective dimensionality of neural activity, N_eff_, as a measure that penalizes small eigenvalues which could arise due to noise. Higher N_eff_ values suggest a high coding dimensionality^37^. A cell with a high temporal dimensionality can have a time-varying magnitude of stimulus selectivity or changes in its order of stimulus selectivity over time (e.g. mixed selectivity). Effective dimensionality during encoding of visual stimuli increased before it plateaued, showing a significantly higher temporal coding dimensionality in mature PFC neurons (F(1, 354) = 3.66, p = 0.000156, GAM; significant developmental improvements -31 to 8 months relative to the mid-adolescence age; Fig. 4a).

In recent years, it has been recognized that population measures of neural activity contain information about stimuli and tasks that may not be apparent in single-neuron measures. Higher dimensionality of population responses, in particular, has been linked to greater cognitive flexibility ^38,39^, which is known to improve in adolescence ^40^. We thus hypothesized that the neural activity followed a high dimensional trajectory not only in single neuron temporal space, but within the full neural state space as well. We therefore calculated the dimensionality of neural responses in the full N-dimensional neural space (where N is the number of neurons in each maturation interval, see methods) across different maturation intervals. Indeed, the dimensionality of the full neural space increased over adolescent maturation (F(1, 9999) = 8079, p <2e-16, GAM; significant increase -31 to -1 months relative to mid-adolescence age; peak at -1 month; Fig. 3b).

The PFC maintains working memory information and manages cognitive processes such as suppressing irrelevant stimuli or distractors during task performance by dynamically representing stimuli in a task-specific manner^41,42^. The same stimuli can be represented in different subspaces when used in the context of different tasks, e.g. creating distinct dimensions for sensory and memory representations to mitigate interference between sensory inputs and memory representations^39,43^ . Since working memory and resistance to distractors becomes increasingly efficient during adolescent development, we speculated that the representation of cue and distractor stimuli may be rotated orthogonally in the neural population space and this relative rotation may improve with maturation, allowing PFC the ability to filter out distractors better. We therefore utilized the distractor task (Fig. 1b), which required animals to view stimuli drawn from an identical set of locations and to keep track of the context of the sensory inputs. In each maturation interval, we applied PCA on a firing rate matrix constructed with each neuron’s average firing rate to each stimulus separately in cue and distractor epochs. The first three principal components generally explained more than 50% of the variance. To ensure fair comparations between different populations of neurons at each maturation interval, we reduced the population firing pattern to the first 3 dimensions to measure the representation structure in low dimensional space. We quantified the differences in representation of the same stimuli based on the angle between subspaces defined by the different task context (target or distractor; Fig. 3c; see methods). The acute angle between the target and distractor subspaces is the largest around mid-adolescence age (peak at -4 month relative to the mid-adolescence age). The representations of the same stimuli during the cue and distractor epochs increased in early adolescence and decreased slightly before plateauing with a significant angle of rotation, indicating that the orthogonal coding emerged during adolescent development (F(1, 7499) = 343.5, p <2e-16, GAM; significant increase -30 to -4 months relative to the mid-adolescence age; Fig. 3d).

**Figure 3.**
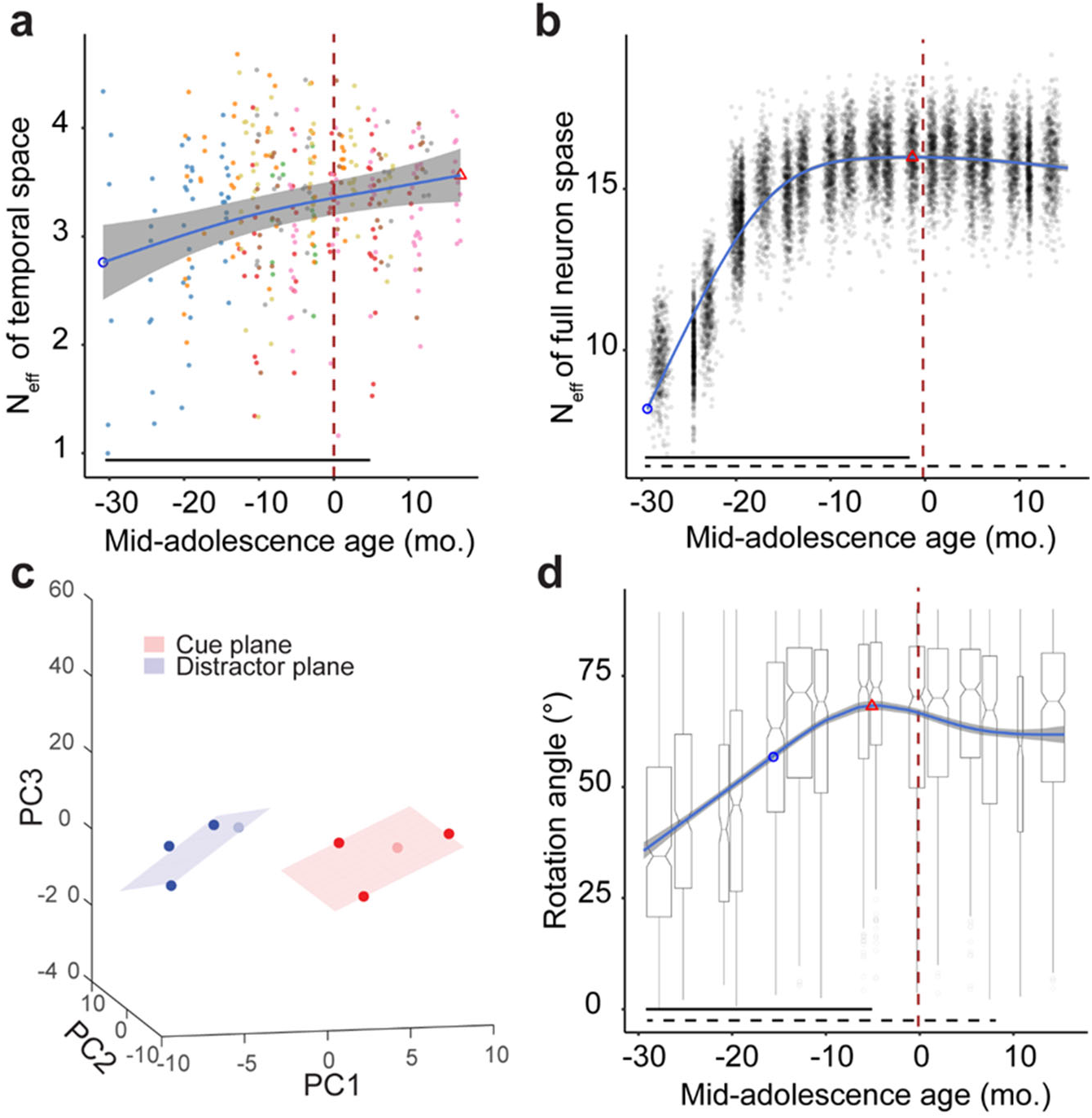
Neural Dimensionality and subspace rotation as a function of mid-adolescence age. (a) Effective temporal dimensionality as a function of mid-adolescence age. Each dot is the N_eff_ estimated in a single neuron. Blue line shows the GAMM fitted trajectory. Gray shaded regions denote the 95% confidence intervals (CIs). Blue circle denotes the time of peak development velocity. Red triangle denotes the time of maximum or minimum value. The dashed vertical line denotes the mid-adolescence age as time 0. The horizontal bar denotes significant developmental effect intervals. (b) Full Neural Dimensionality. Scatter plots show distribution of the dimensionality of the full neural response during the stimulus presentation and delay epochs. Each dot is the estimated effective dimensionality of a pseudo population of neurons at the average mid-adolescence age of the neurons (see methods). Blue line showed the GAMM fitted trajectory. Gray shaded regions denote the 95% confidence intervals (CIs). Blue circle denotes the time of peak development velocity. The red triangle denotes the time of maximum dimensionality. The dashed vertical line denotes the mid-adolescence age 0. The horizontal bar denotes significant developmental effect intervals. (c) An example of low dimensional representation of spatial stimuli during the cue and distractor epoch in correct trials in ODR with distractor task (population mid-adolescence age = -20 month). Rotation φ=44°. (d) Rotation angle between cue and distractor subspace as a function of mid-adolescence age. Each dot is the rotation angle of a pseudo population of neurons at the average mid-adolescence age of the neurons (see methods). Blue line showed the GAMM fitted trajectory. Gray shaded regions denote the 95% confidence intervals (CIs). Blue circle denotes the time of peak development velocity. The red triangle denotes the time of maximum rotation angle. The dashed vertical line denotes the mid-adolescence age 0. The horizontal bar denotes significant developmental effect intervals.

### Some structural brain changes predict adolescent cognitive improvements

We wished to determine the aspects of brain structural changes that best explain changes in neural activity and behavior we observed. Since neurophysiological recordings were obtained from these animals, we performed volume measurements of cortical and subcortical structures on the hemisphere opposite to the one where recordings were performed. We first considered global brain measures for white and gray matter (Extended Data Fig. 4-10), subcortical areas (Extended Data Fig. 11-12) and fiber tracts (Extended Data Fig. 14-16). Overall, we observed increases in all brain structure volumes (Fig. 4a-c). We also examined more specific measures in brain areas implicated with working memory, including the lateral prefrontal cortex (Fig. 5).

In humans, gray matter (cortical) volume peaks during childhood and decreases thereafter ^3^, thought to be driven by processes such as synaptic pruning. In monkeys, a global volume maximum is also observed in childhood (0.74 years), however many areas exhibit a second period of increase in adolescence ^4^. Indeed, in our cohort, which was tracked after year 3, we observed increases in all brain structure volumes of the macaque’s brain (Fig. 4a-c). We found that whole brain cortical matter volume increased in our dataset (F(1, 75) = 4.2, p = 0.0467, GAM; significant increase -33 to - 12 months relative to the mid-adolescence age; Fig. 4b). Peak gray matter volume was observed at a chronological age of 78 months, in late adolescence. Although volume increase was relatively small in the cerebral cortex (3.5% from -43 to 20 months relative to the mid-adolescence age, Fig. 4b), it was more pronounced in non-cortical areas and particularly white matter (15.3%, Fig. 4b), subcortical structures (10.0%, Fig. 4b), diencephalon (20.0%, Fig. 4c), mesencephalon (22.0%, Fig. 4c), myelencephalon (39.0%, Fig. 4c), cerebellum (12.5%, Extended Data Fig. 10b), and cerebellum white matter (21.6%, Extended Data Fig. 10c). Cerebral white matter exhibited a relatively linear increase in adolescent monkeys (F(1, 75) = 25.8, p = 2.5e-6, GAM; significant increase -43 to 20 months relative to the mid-adolescence age; Fig. 4a-b), as observed in humans.

As expected, the variation in lobar cortical volumes (Fig. 4g), thickness (Fig. 4h), and surface area (Fig. 4i) was minimal, corresponding to the relatively small variation in whole brain cortical volumes. However, the variation and the inverse U-shape of the frontal lobe cortical thickness stands out when compared to metrics or other lobes (Fig. 4e). To investigate this further, we focused on the finer segmentation of brain regions and fiber tracts associated with working memory and the frontal cortex (see Methods - Fig. 5a-f, Extended Data Fig. 6). Thickness in lateral PFC and motor cortex increased early on and peaked before the mid-adolescence age, followed by a decrease later in adolescence (Fig. 5b, e). This was also the case in posterior parietal cortex, a region functionally and anatomically highly interconnected with the lateral prefrontal cortex (Extended Data Fig. 5d, 7). In contrast, most other frontal cortical regions including the orbitofrontal cortex and anterior cingulate cortex did not show any significant change in cortical thickness with maturation (Fig. 5b, e). Most regions in the other lobes (parietal, temporal, occipital) did not show significant variations in the three metrics, except for primary areas such as the somatosensory cortex (Extended Data Fig. 4) and the primary visual cortex (Extended Data Fig. 5), as well as the previously mentioned motor cortex (Fig. 5b, e).

Even for the case of the lateral prefrontal cortex, the volume of which exhibited an overall decrease, volume did not predict behavioral performance well (DI vs. volume: r = 0.132, permutation test, p = 0.394; RT vs. volume: r = -0.171, p = 0.1800). Thickness and surface area were only weakly correlated with measures of behavioral performance (DI vs. thickness: r = 0.031, p = 0.963; RT vs. thickness: r = -0.299, p = 0.008; DI vs. surface: r = 0.422, p < 0.0001; RT vs. surface: r = 0.106, p = 0.569).

**Figure 4.**
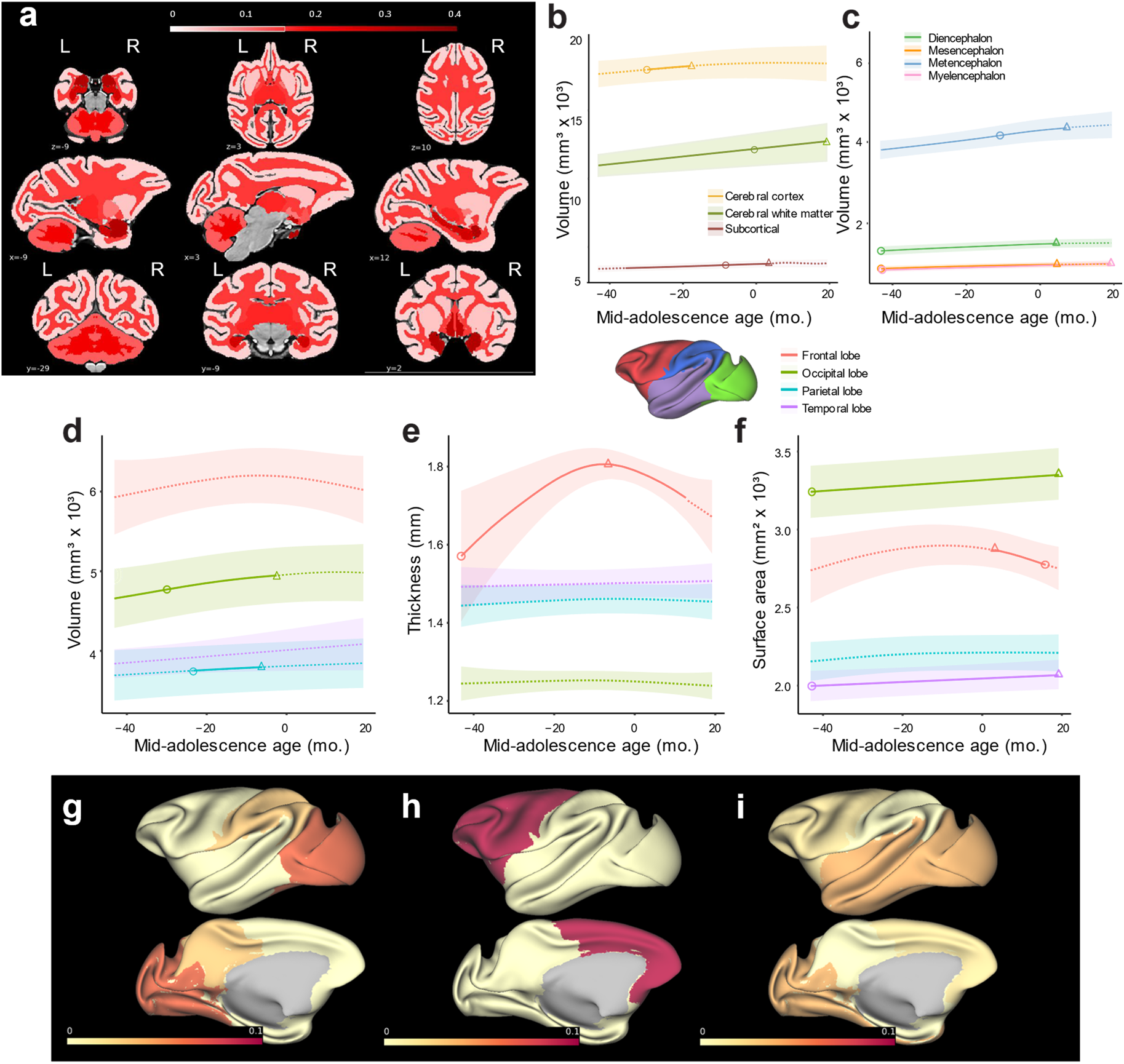
Longitudinal developmental trajectories of the macaque brain. (a) Map of volume changes for white and gray matter and subcortical areas. (b and c) Volumes of brain segmentations and structures as a function of age. (d-f) In each lobe, segmented using Charm atlas level 1, (d) volumes, (e) thickness, (f) surface as a function of age. Solid curves indicate significant developmental effect of the GAMM fittings. Shaded ribbons denote the 95% confidence intervals (CIs). Circle denotes the time of peak development velocity. Triangle denotes the time of maximum value. (g-i) Proportional change during adolescence compared of different lobes for (g) volume, (h) cortical thickness and (i) surface area. Color represented proportional change of each lobe relative to the earliest data points (-43 months to the mid-adolescence age).

Lastly, developmental effects on white matter maturation, quantified using fractional anisotropy (FA), radial diffusivity (RD) and mean diffusivity (MD), were evident across the brain, with most tracts reflecting increases in FA and decrease in RD and MD with development (Fig. 5g, Extended Data Fig. 13). We also examined in more detail tract maturation that most closely paralleled the trajectory of working memory performance improvement and prefrontal cortical activity changes. We performed correlation analyses between tracts we identified and behavioral performance of ODR task including DI and RT. FA changes in several white matter tracts had high negative correlation (|r| > 0.9) with the precision (DI) of behavioral performance in the ODR task (Extended Data Fig. 18). These tracts include the middle longitudinal fasciculus (MLF), superior fronto-occipital fasciculus (SFOF), and anterior cingulum which projected from or to the frontal lobe (permutation test, p<0.0001, in each case). Strong negative correlations were also observed between FA trajectories with RT during adolescence (Extended Data Fig. 19).

**Figure 5.**
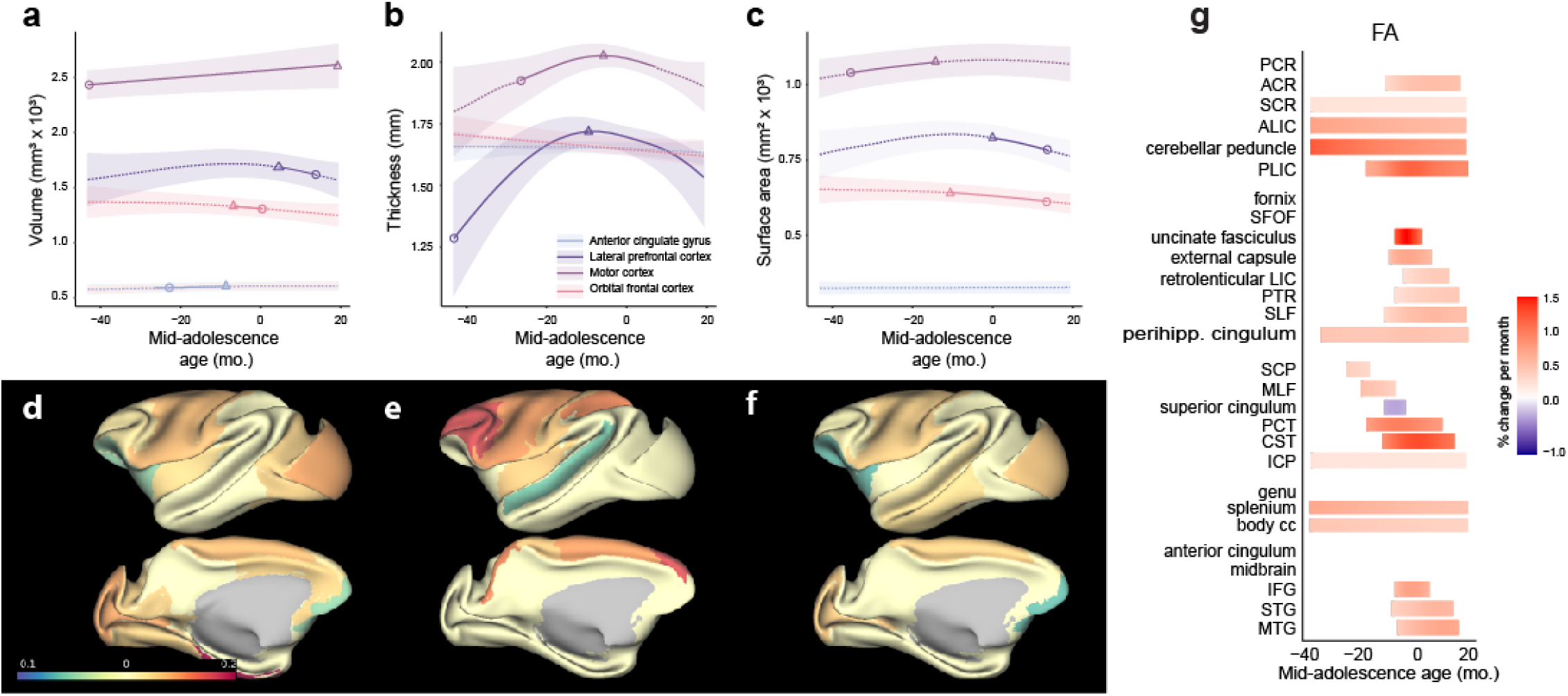
Structural changes during development in frontal lobe and white matter tracts. (a-c) Longitudinal developmental trajectories of areas in the frontal lobe for (a) volume, (b) cortical thickness and (c) surface area. Solid curves indicate significant developmental effect of the GAMM fittings. Shaded ribbons denote the 95% confidence intervals (CIs). Circle denotes the time of peak development velocity. Triangle denotes the time of maximum volume. (d-f) Proportional change during adolescence compared of different brain areas in Charm atlas level 2 for (d) volume, (e) cortical thickness and (f) surface area. Color represents proportional change of each lobe relative to the earliest data points (-43 months to the mid-adolescence age). (g) Fractional anisotropy (FA) of major white matter tracts as a function of age. Each row is an ROI grouped to projection, association, commissural tracts, brainstem white matter regions and short-range white matter. Rows are sorted by time of maturation in each group.

## Discussion

Human cognitive ability improves significantly from childhood through adolescence ^7,36^, but the underlying changes in neural activity have not been hitherto well investigated. In this study, we illuminate the processes underlying cognitive maturation, employing non-human primates as a model. Monkeys, as humans, exhibited substantial variability in development, we therefore relied on skeletal assessment to best physical development and align growth trajectories of different individuals. We used an ostensibly simple task to track the trajectory of working memory function, using the precision and latency of responses in the oculomotor delayed response task as primary behavioral outcomes. This choice allowed us to track performance from the earliest time points, as all animals were able to master that readily and to compare results with the extensive human childhood and adolescence literature ^7,8^. Critically, recent studies have shown that these variables track adolescent development remarkably well across a range of abilities, including much more complex processes such as planning, response inhibition, and other aspects of executive function^10^. Neuropsychiatric conditions and mental illnesses that emerge in adolescence to early adulthood, most notably schizophrenia, are also associated with decreased performance in working memory tasks ^44^. Our results revealed a progressive improvement in both precision and latency (defeating a speed-accuracy tradeoff) with a shape that closely resembled human development.

Neurophysiological recordings from the prefrontal cortex revealed the changes in neural activity that could account for these behavioral changes. Prior studies had suggested higher firing rates in adult animals particularly in the delay period of working memory tasks ^23^. Our current results suggested that changes even in the baseline firing rate of prefrontal neurons during the fixation period of the task could capture well the trajectory of behavioral improvement. Computational models suggest that recurrent connections between prefrontal neurons allow activity to reverberate in the network during working memory tasks, and increased synaptic drive will generally render a network more stable ^45^ ^46^. The improvement in saccade precision metrics indicates that decrease in trial-to-trial variability is an important developmental milestone. Indeed fMRI measures related to fluctuations in the amplitude of task-related brain states stabilize during development ^47^. Excessive variability is also a sensitive marker of cognitive function in conditions such as schizophrenia ^48^ and ADHD^49^. We now tie these behavioral variability measures to changes in neural variability metrics, including the coefficient of variation of inter-spike intervals and Fano factor. Reduced behavioral and firing rate variability, too, can be mapped to stronger synaptic drive in neural networks, which renders the peak of activation in the network less susceptible to drift in each trial ^50^ . Variability may also reflect a period of plasticity in adolescence as circuits are being engaged differently to identify optimal function into adulthood in a Hebbian process selecting circuitry that is most highly used due to its efficacy.

Our analysis also revealed more complex measures of neural activity that change during adolescent development. The dimensionality of neural representations increased, as did the extent of rotation of stimulus representations when these were presented as cue stimuli, or as distractors, in a variant of the ODR task. Higher dimensionality of population responses, and the related phenomenon of nonlinear mixed selectivity has been linked to greater cognitive flexibility ^38,51^. Rotation of stimulus representation is a critical mechanism in suppressing irrelevant stimuli or distractors during task performance by dynamically representing stimuli in a task-specific manner^39,41–43^ . Such changes observed during development now account for cognitive improvements in domains such as cognitive flexibility^52^ and ability to withstand distraction^22^, which improve through adolescence.

Recent structural imaging studies have codified the trajectory of age-related brain changes across cohorts of hundreds of individual humans ^3^ and monkeys ^4^. A hallmark of brain development is the decrease of cortical volume and thickness after childhood ^5^, thought to be driven by pruning of infrequently used synapses ^12^, although the apparent decrease in thickness may also be driven by myelination of gray matter fibers ^53^. Pruning confers clear improvements in neural computations ^54^, however decreases in gray matter volume were not drivers of the working memory enhancement that we observed. Instead, our findings pointed out an association between the maturation of specific white matter tracts, and improvements in working memory performance and prefrontal cortical activity during adolescence. These results align with existing literature that highlights the prolonged maturation of white matter pathways in humans, which continues well into early adulthood ^55,56^. The observed FA increases in tracts connecting the frontal lobe to cortical and subcortical regions are consistent with findings that show an association between the maturation of white matter and improvements in cognitive tasks that depend on these connections, such as working memory and executive functions. Taken together, these results show new evidence for significant neural maturation through adolescence into adulthood. Disruptions in the maturation of these systems may play an important role in psychopathology that emerges at this time (e.g., schizophrenia, mood disorders, substance use disorders) and are typically associated with impairments in executive function.

## Methods

### Subjects

Behavioral, imaging, and neurophysiological recordings were obtained from a total of 14 (10 male and 4 female) rhesus monkeys (*Macaca mulatta*) of three cohorts: Cohort A: eight monkeys (6 male, 2 female); cohort B :four monkeys (4 male); and cohort C: two monkeys (2 female). All surgical and animal use procedures were reviewed and approved by the Institutional Animal Care and Use Committees of Wake Forest University and Vanderbilt University, in accordance with the U.S. Public Health Service Policy on Humane Care and Use of Laboratory Animals and the National Research Council’s Guide for the Care and Use of Laboratory Animals.

### Developmental profiles

We tracked developmental measures of Cohort A monkeys on a quarterly basis before, during, and after neurophysiological recordings. Similar to previous studies ^23,24^, we obtained morphometric measures including body weight, trunk length, femur length, canine eruptions and length. Testicle length, width and volume was additionally determined for male monkeys (with a Prader Orchidometer, ESP Limited, Rustington, UK), and nipple length was determined for female monkeys. To determine each monkey’s developmental progress, we determined bone maturation via X-rays of upper and lower extremities, and assayed hair concentration of hormones including testosterone and dihydrotestosterone. We relied primarily on skeletal assessment to best capture physical development ^35,57,58^ and align the growth trajectories of different individuals. Using these measures, we defined a mid-adolescence age for each monkey, defined as the time of each monkey’s distal tibial epiphyseal closure, as observed by veterinary professionals evaluating the X-rays, blind to findings of other aspects of the study.

### Behavioral Tasks

Monkeys were trained to perform an oculomotor delayed response (ODR) Task ^59^. This is a spatial working memory task that requires the subject to remember the location of a cue stimulus appearing on a screen for 0.5 s (Fig. 1a). The cue stimulus was a 1° white square appearing at one of eight locations arranged on a circle of 10° eccentricity, spaced by 45 degrees (Fig. 1c). After a 1.5 s delay period, the fixation point was extinguished, indicating the monkey to make a saccade to the remembered location of the cue within 0.6 s. The saccade needed to terminate on a 6° radius window centered on the stimulus, and the monkey was required to hold fixation within this window for at least 0.1 s. Animals were rewarded with liquid rewards (typically fruit juice) for successful completion of a trial.

Eye position was monitored with an infrared eye tracking system (sampling rate 240 Hz, ISCAN, RK-716; ISCAN, Burlington, MA). Breaking fixation at any point before the offset of the fixation point aborted the trial and resulted in no reward. The visual stimulus display, monitoring of eye position, and synchronization of stimuli with neurophysiological data were performed using in-house software, and implemented with MATLAB ^60^.

Four of the eight monkeys of cohort A (one female and three males) were additionally trained on the ODR task with a longer delay period of 3.0 s after they were capable of performing the shorter task. The same four monkeys were also trained to perform a variant of the task, the ODR with Distractor Task ^61^. This task involved a distractor, presented 1.25 s after the cue for 0.5 s, followed by another 1.25s delay period before fixation point was off, making the total duration of the trial the same as in the ODR task with 3 s delay. The cue could appear at one of four locations arranged on a circle of 10° eccentricity, spaced by 90 degrees. For each cue location, five distractor conditions were used: the location of the distractor could appear at location that is diametric to the cue (180 degree), 90 degrees counterclockwise to the cue (90 degree), 45 degrees counterclockwise to the cue (45 degree), or the same as the cue (0 degree). The fifth condition involved no distractor presentation.

### Recording Phases

The monkeys were initially trained in the tasks mentioned above before their neurophysiological recordings. They were naïve to behavioral training or task execution of any kind prior to the behavioral training. Once the animals had reached asymptotic performance, behavioral and neurophysiological recordings were obtained from the monkeys of cohort A at time points spaced approximately 3 months apart from 3.4 to 6.2 years old. Each behavioral time point of each animal contained an average of 19 sessions. Between time points, the animals were returned to their colony and were not tested or trained in any task until their next time point.

### MRI acquisition and processing

Structural MRIs were collected from the monkeys of cohort A every 3 months from 2.8 years (34 months) of age to 5.8 years (69 months) of age. In preparation for the MRI scan, anesthesia was induced using ketamine (5–10mg/kg) and dexmedetomidine (0.015mg/kg), and was maintained using isoflurane. The animals were intubated and artificially ventilated at about 20 breaths per minute. Expired CO_2_ was monitored and maintained between 35 and 45 mmHg. Animals were scanned under isoflurane anesthesia at 1%–1.5%. Heart rate and oxygen saturation levels were monitored using a pulse oximeter. Their body temperature was maintained using warm blankets. The MRI system was a 3 Tesla Siemens MAGNETOM Skyra (Siemens Healthcare, Erlangen, Germany). Anatomical images were acquired using a T1-weighted MPRAGE sequence: TR = 2700 ms, TE = 3.32 ms, inversion time = 880, FOV = 128 × 128 mm, 192 slices of 0.5 mm thickness, resolution = 0.5 mm isotropic. Resting state time series data were also acquired using a multiband EPI sequence: TR = 700 ms, TE = 32.0 ms, flip angle = 52°, repetitions = 700, FOV = 128 × 128 mm, 32 slices, resolution = 2 mm isotropic.

Spatial pre-processing was performed using a pipeline coded in python, which relied on functions from AFNI,^62^ ANTs^63^, FSL^64^, FreeFurfer^65^ and Connectome Workbench^66^ for inhomogeneity correction, spatial and surface registration to a standardized space, the study template space. The study template was based on the average of the last T1 anatomical image (T1^last^) of each animal when registered in a common space using the function “anats_to_common” from Sammba-MRI^67^. A high-resolution NMT template (NIH Macaque Template) as well as the CHARM^68^ and SARM^69^atlases segmentation were registered to the study template. The segmentation atlas was created directly in the NMT template space using automated and manual segmentation (Fig. 4a, 5a). Individual anatomical T1 images (T1^n^) were registered to their T1^last^ and each T1^last^ was registered to the study template. The two movement parameters (T1^n^ to T1^last^ and T1^last^ to study template) were combined to register T1^n^ to the study template. Inversion of these movement parameters was used to register the atlases to the individual T1^n^ images. The co-registrations of the atlases to T1 were individually inspected and corrected manually when necessary. The volumes (in mm^3^) were calculated using the function “3dhistog” from AFNI. Surface and thickness were calculated using the Connectome Workbench. Figures were produced using the Connectome Workbench and nilearn^70^.

### DTI preprocessing

DTI data were acquired in pairs with a reversed phase encoding direction in the second scan (e.g., PA vs AP). A diffusion-weighted spin-echo echo-planar imaging sequence was utilized to obtain 82 whole-brain slices of 2mm thickness in 30 directions. Data were processed for analysis using MRtrix3 ^71^ and the Oxford Centre of fMRI of the Brain Software Library (FSL). The raw DICOM images acquired from the scanner were converted to NIFTI format using dcm2nii, and the corresponding bval and bvec files containing information pertinent to the diffusion gradient were combined across scans. Images were then denoised (“dwidenoise”) and mean b=0 images were calculated.

Following this, susceptibility induced and eddy current distortion was corrected using FSL (“TOPUP”) and (“eddycorrect”) respectively. A tensor model was fitted to each voxel (“dwi2tensor”) and fractional anisotropy (FA) maps were calculated (“tensor2metric”). A mask was also created from the T1-weighted image (“bet”), segmenting the brain and non-brain tissue from the whole head.

Each FA image was then registered to the subject’s respective skullstripped T1-weighted image by affine transformation (“flirt”). These co-registered images were subsequently registered to a diffusion-tensor-based white matter atlas for rhesus macaques ^72^. This allowed for a group analysis of several parameters: 1) the directionality of water diffusion within white matter tissue (FA), 2) the mean apparent diffusion coefficient of the diffusion tensor (mean diffusivity (MD)), 3) the principal eigenvalue, or diffusion parallel to the principal axis of diffusion (axial diffusivity -AD), and 4) the mean of the two non-principal eigenvalues, or the diffusion perpendicular to the principal axis of diffusion (radial diffusivity - RD). The MD, AD, and RD maps were derived from the diffusion tensor using the same method employed for generating the FA maps.

After initial processing, a whole-brain region-of-interest (ROI) analysis was performed, with 53 white matter tracts from the DTI atlas selected. Left and right hemispheres were separately calculated and averaged together.

### Surgery and neurophysiology

After the animals of cohort A had reached asymptotic performance in the behavioral tasks for the first time, we implanted a 20-mm diameter recording cylinder over the prefrontal cortex of each monkey. Localization of the recording cylinder and of electrode penetrations within the cylinder was based on MR imaging, processed with the BrainSight system (Rogue Research, Montreal, Canada). Recordings in each time point (see recording phases) were collected with glass or epoxylite coated Tungsten electrodes with a diameter of 250 μm and an impedance of 4 MΩ at 1 KHz (FHC Bowdoin, ME). Electrical signals recorded from the brain were amplified, band-pass filtered between 500 Hz and 8 kHz, and stored through a modular data acquisition system at 25 μs resolution (APM system, FHC, Bowdoin, ME).

Recordings were obtained and analyzed from areas 8a and 46 of the dorsolateral prefrontal cortices. Neurons were not pre-screened prior to collection; we recorded from all neurons isolated from our electrodes. Recorded spike waveforms were sorted into separate units using a semi-automated cluster analysis method based on the KlustaKwik algorithm ^73^. Neurons for which at least 4 correct trials in every stimulus condition were available in the ODR task were used in the following analyses.

### Behavioral analyses

We analyzed task performance in the ODR task and distractor task as the percentage of trials that resulted in correct responses and by determining the animals’ saccade precision and reaction time. For saccade precision analysis, we calculated each session’s saccade Dispersion Index (DI), defined as the area within one standard deviation from the average landing position of each target condition ^74^ . We calculated reaction times by determining the interval between the offset of the fixation point and the time of saccade onset.

### Firing metrics

We measured three metrics of neuronal firing from different task epochs using correct trials in the ODR tasks: firing rate, Fano factor, and coefficient of variation (CV) of firing rate and interspike intervals (ISI), to characterize the intrinsic firing pattern for neuron at recorded at different maturation stages. The firing rate measures the activity of the neurons and the Fano factor and CVs measure the variability of the neural activities. The CV of ISI is defined as:

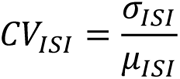

where σ*_ISI_* is the standard deviation of interspike intervals and *μ_ISI_* is the mean of the interspike intervals. Similarly, the CV of firing rate is defined as:

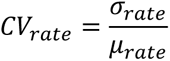

where *σ_rate_* is the standard deviation of firing rate during baseline across all included trials and *μr_ate_* is the mean of the baseline firing rate. Fano factor was computed as:

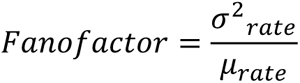

We computed the Fano factor of spike counts to measure the variance of the firing rate as in previous studies ^75^. In brief, data for each condition in each neuron are initially treated separately. For each condition, we computed the variance (across trials) and mean of the firing rate during the epoch of interest. The Fano factor was the slope of the regression relating the variance to the mean.

### Single neuron tuning

To investigate single neuron tuning, we used firing rates from eight classes and fitted a Gaussian curve to the data. We defined the locations and corresponding firing rates. The Gaussian model function used to describe the tuning curve was:

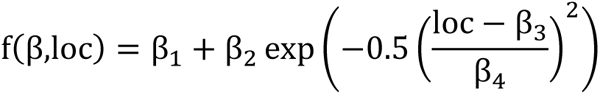

where: *β*_1_ is the baseline firing rate, *β*_2_ is the amplitude of the Gaussian peak, *β*_3_ is the location of the peak, and *β*_4_ is the standard deviation (width) of the Gaussian curve. The lsqcurvefit function in MATLAB was used to fit the Gaussian model.

### Intrinsic Timescales

To calculate the intrinsic timescale of single neurons, we constructed a data matrix where each row represented a trial and each column represented time bins for a single neuron, with the bin size of 50 ms. We used the 1000 ms pre-cue fixation epochs of each neuron. We excluded neurons with an average firing rate of less than 1 spike per second and those with any time bins containing zero spikes across all trials. Neurons with poor fitting quality (e.g., intrinsic time scales greater than 0.5 seconds) were excluded from the final analysis as well ^76,77^. The average autocorrelation was calculated for each time lag and normalized by the total variance of the spike counts. We then fitted an exponential decay model to the autocorrelation data using:

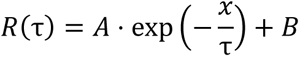

where *A* is the amplitude, *τ* is the intrinsic time scale, and *B* is an offset parameter. The fit function in MATLAB was used to fit the exponential model.

### Information contained in single neurons

We used the bias-corrected percentage of variance-explained (ωPEV) statistic to estimate single neuron selectivity to task conditions, i.e. locations of visual stimuli ^78^. We calculated ωPEV using the measures of effect size (MES) toolbox of MATLAB, version 1.6.0.0, which calculate the amount of variance in the neurons’ firing rate explained by the location of the stimuli. ωPEV is defined as:

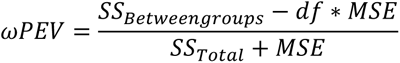

where *SS_Total_* denotes the total sum of squares, i.e., total variance, *SS_BetweenGroups_* denotes the variance between groups, df the degrees of freedom, and MSE the mean-squared error.

### Temporal coding dimensionality

Firing metrics use averaged measures in certain time windows and do not reflect the consistency or dynamic of neural responses over time. Therefore, we used principal component analysis (PCA) to quantify temporal coding stability for each individual neuron. We estimated the effective dimensionality (N_eff_) of the temporal state space of each neuron ^37^. PCA was performed on the mean firing rate of trials in the same task condition across different time bins during the cue presentation, thereby quantifying the temporal variability of task-dependent firing. For each neuron, we then constructed a data matrix comprising n trials × 10 time bins. For each trial, we computed the firing rate in each time bin. Each time bin is a 50ms time in the 500ms stimulus presentation. Then, we averaged binned firing rates by task condition (8 locations). Columns in the matrix correspond to 10 adjacent, independent time windows spanning 500 ms and rows to the 8 task condition averages. PCA is then performed on the matrix to quantify and arrange the variance along the principal components in the subspace spanned by the ten independent time bins. The eigenvalues associated with each principal component give us the means to quantify the effective dimensionality (N_eff_) of our temporal state space:

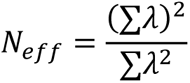

where λ represents the eigenvalues.

### Dimensionality in full neural space across maturation intervals

Similar to dimensionality in temporal space of each neuron, we calculated the dimensionality of neural responses in the full N-dimensional neural space (where N is the number of neurons), to test global changes in the neural space across maturation. For the population analysis, neurons from all animals were pooled and analyzed together. We segmented the neurons based on their mid-adolescence age into 20 even time intervals (each ∼200 days long), spanning from the earliest to the latest session recorded across all animals. For each maturation interval, we combined neurons across animals to create a pseudo-population. We averaged the firing rate per condition (8 locations in ODR task) within 200ms bins, stepped by 100ms. The average response was calculated for each condition in each time bin during the delay period of the task (1.5 s). To control for the effect of sample size on dimensionality, we randomly sampled 30 neurons from each pseudo-population, to ensure no oversampling from any populations in each iteration. PCA was performed on the concatenated data: size = (condition × time) × 30 neurons. We then calculated the effective dimensionality (N_eff_) of each maturation interval. The distribution of N_eff_ values was estimated with a bootstrap: we resampled neurons with replacement per maturation interval for 500 times and then performed the estimation of N_eff_. This gave us a dataset of 10000 values of N_eff_ and these values were used to estimate the change in N_eff_ across adolescent development and fitted with GAMM.

### Rotation of subspaces

We applied PCA to denoise and quantify the difference between representation of the target and distractor. PCA was performed on the mean firing rate of neurons across 15 maturation intervals (200 days) from 4 animals who performed the distractor task.

For each neuron, we calculated the average firing rate of the cue and distractor epochs in all correct trials for different task conditions (4 locations). The firing rates of each neuron formed an 8×1 vector (4 locations × 2 epochs). The population activity matrix sizes 8 × N where N is the number of neurons in the population used. Then we z-scored the population activity matrix to account for the baseline firing difference among neurons. To find the rotation between two periods, we applied PCA on the z-scored population matrix. We selected the first three eigenvectors of the covariance and sorted them by decreasing order in terms of explaining the variance.

We then projected the population activity matrix into a three-dimensional (3D) PCA space. The population representations of each of the four spatial locations in two task epochs was represented by a vector. Four spatial locations in each of two task epochs then formed a subspace respectively spinning in the full space. We found the best-fit hyperplanes for cue and distractor subspaces by least squares to minimize the distance from each point to the hyperplane, implemented using fminsearch function in MATLAB. To examine the relationship between the two subspaces, the rotation is defined as the acute angle between the normal vectors of the two best fit planes.

Similar to effective dimensionality, we randomly sampled 30 neurons from each of the 15 intervals to control the effect of sample size. The distribution of rotation angles was estimated with a bootstrap: we resampled neurons with replacement per maturation interval 500 times and calculated the rotation angle in each resampling. The total dataset contained 7500 values of rotation angles and was then used to estimate the change in rotation of subspaces across adolescent development and fitted with GAMM.

### Generalized additive mixed models

We sought to characterize the adolescent development of non-human primates in cognitive function, prefrontal cortex activity and the brain regions and structures that support in our longitudinal sample. We expected developmental stages to exhibit a nonlinear relationship with each outcome. Therefore, we utilized generalized additive mixed models (GAMM). GAMM is a flexible, semiparametric method for identifying and estimating nonlinear effects of covariates on the outcome variable when observations are not independent ^79^.

All GAMM analyses were implemented using the mgcv package for R to fit a series of GAMMs for the outcomes of interest, each with a smooth function of mid-adolescence age as a covariate, using a thin plate regression spline basis to estimate this smooth function. Random effects in each GAMM included subject-specific intercepts and slopes for mid-adolescence age. For proportional outcomes like behavioral task performance, the quasi-binomial family and logit link function was used. For all other outcomes, which are continuous, a Gaussian family and identity link function was used. For those models for which there was a statistically significant fixed effect, the gratia package for R^80^ was used to conduct exploratory post-hoc analyses to identify significant periods of developmental change. Specifically, the derivatives of each estimated smooth function of age were approximated using the method of finite differences, and a simultaneous 95% confidence.

### Correlation between trajectories

To evaluate similarity between developmental trajectories, we calculated the correlation between the GAMM predictions of different measures. To ensure consistency and comparability, the predictor values (mid-adolescence age) were evenly sampled at 100 intervals between the earliest and latest time points for behavioral data. Each curve was then normalized using z-score normalization.

We calculated two key metrics: the Pearson correlation coefficient (r) and the Root Mean Square Error (RMSE). Before calculating RMSE, each normalized curve was shifted such that the starting point was aligned to zero. This was done by subtracting the starting value of the curve. After shifting, the absolute values of the data points were taken to ensure uniformity in the direction of both curves, facilitating a comparable visualization using RMSE between positively and negatively correlated pair of curves. The RMSE was calculated as:

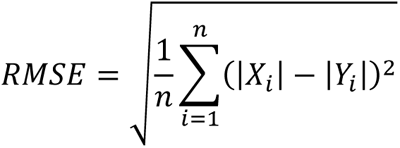

where ∣ *Xi* ∣ and ∣ *Yi* ∣ are the absolute values of the shifted and normalized data points from the respective curves.

We conducted a permutation test with a maxT approach to assess the statistical significance of the observed correlation coefficients. We randomly permuted *X* and *Y* to recalculate the correlation coefficient across 1,000 permutations to simulate the null hypothesis of no correlation. The maxT p-value was calculated to determine the probability of observing a correlation as extreme as the detected one, or more extreme, under the null hypothesis.

## Data availability

Data for the current study will be made available upon publication.

## Code availability

The code used to process the results and generate the figures as well as the pipeline to process the MRI anatomical images will be made available upon publication.

## Acknowledgements

We wish to thank Austin Lodish, Leonardo Silenzi, Jasmine Sun, Gracie Hilber, Chrissy Suell, Katelyn Clemencich, and Susan Apt for technical help; Simon Clavagnier for his support on the processing pipeline development and segmentation of the macaque brain; and Ueli Rutishauser for helpful comments on the manuscript. This work was supported by NIH grants R01 MH117996 and R01 MH116675.

## Author contributions

C. C., E. S., T. R. S, C. T. W., B. L. conceptualized the project and designed the experiments. J. Z., C. M. G., M. X. Z., Z. W. and X. Q. performed experiments. A. M., A. W. A., F. J. C., S. B. H. contributed to the processing of imaging data. J. Z. and C. M. G. performed the analysis.

---

**Extended Data Fig. 1.**
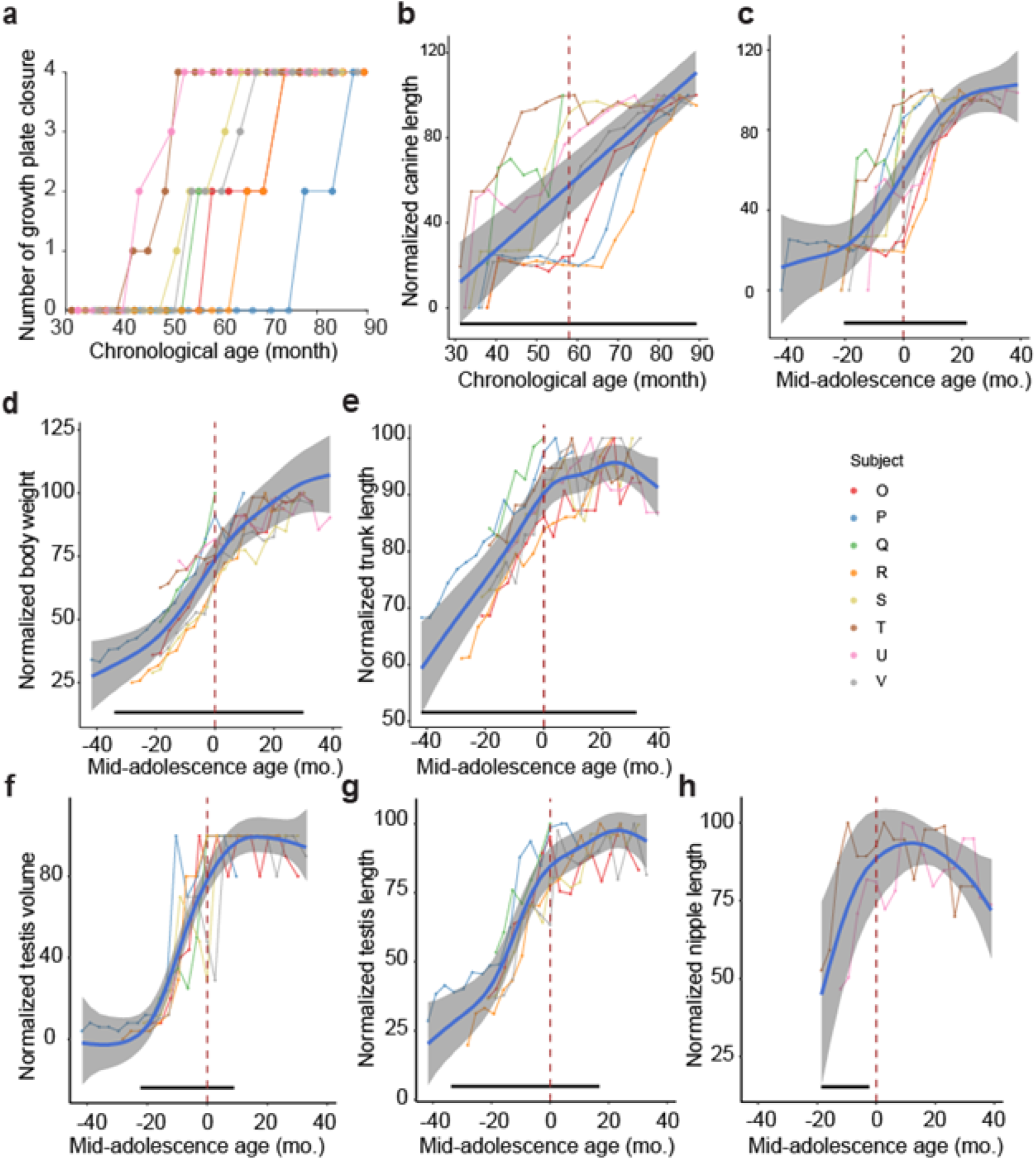
Developmental profile and mid-adolescence age. (a) number of growth plate closure of each monkey as a function of chronological age. (b) Average canine length of the same subjects, aligned on their chronological age. Blue line showed the GAMM fitted trajectory. Gray shaded regions denote the 95% confidence intervals (CIs). (c) same as in (b) but aligned on their mid-adolescence age. (d and e) Body weight and trunk length, presented in the same fashion as in (c). (f and g) Testicle volume and testicle size (length of longer dimension) of the male monkeys, presented in the same fashion as in (c). (h) Nipple length of the female monkeys, presented in the same fashion as in (c). The dashed vertical line denotes the mid-adolescence age 0. The horizontal bar denotes significant developmental effect intervals.

**Extended Data Fig. 2.**
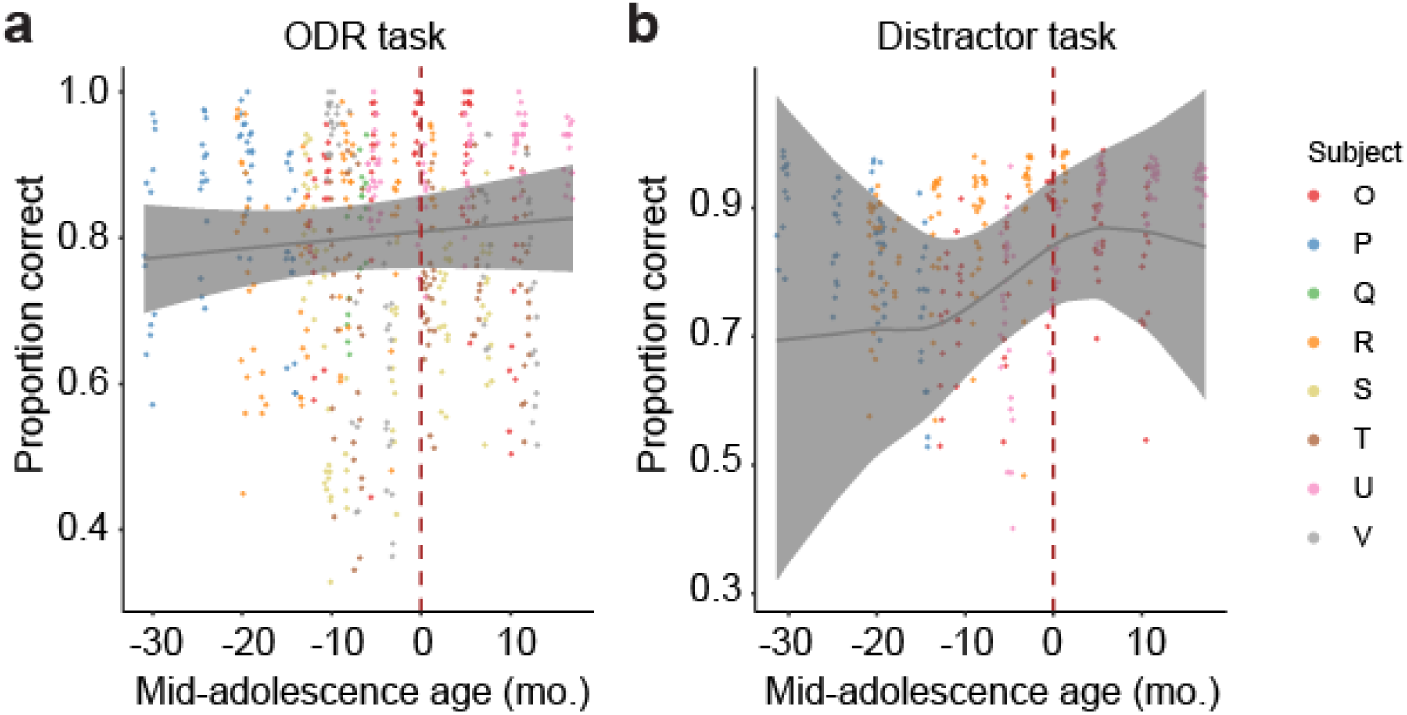
Trial completion performance. Percentage of trials that were deemed correct and rewarded as a function of age. (a) Performance in the oculomotor delayed response (ODR) task. (b) Performance in the ODR with distractor task. The dashed vertical line denotes the mid-adolescence age 0.

**Extended Data Fig. 3.**
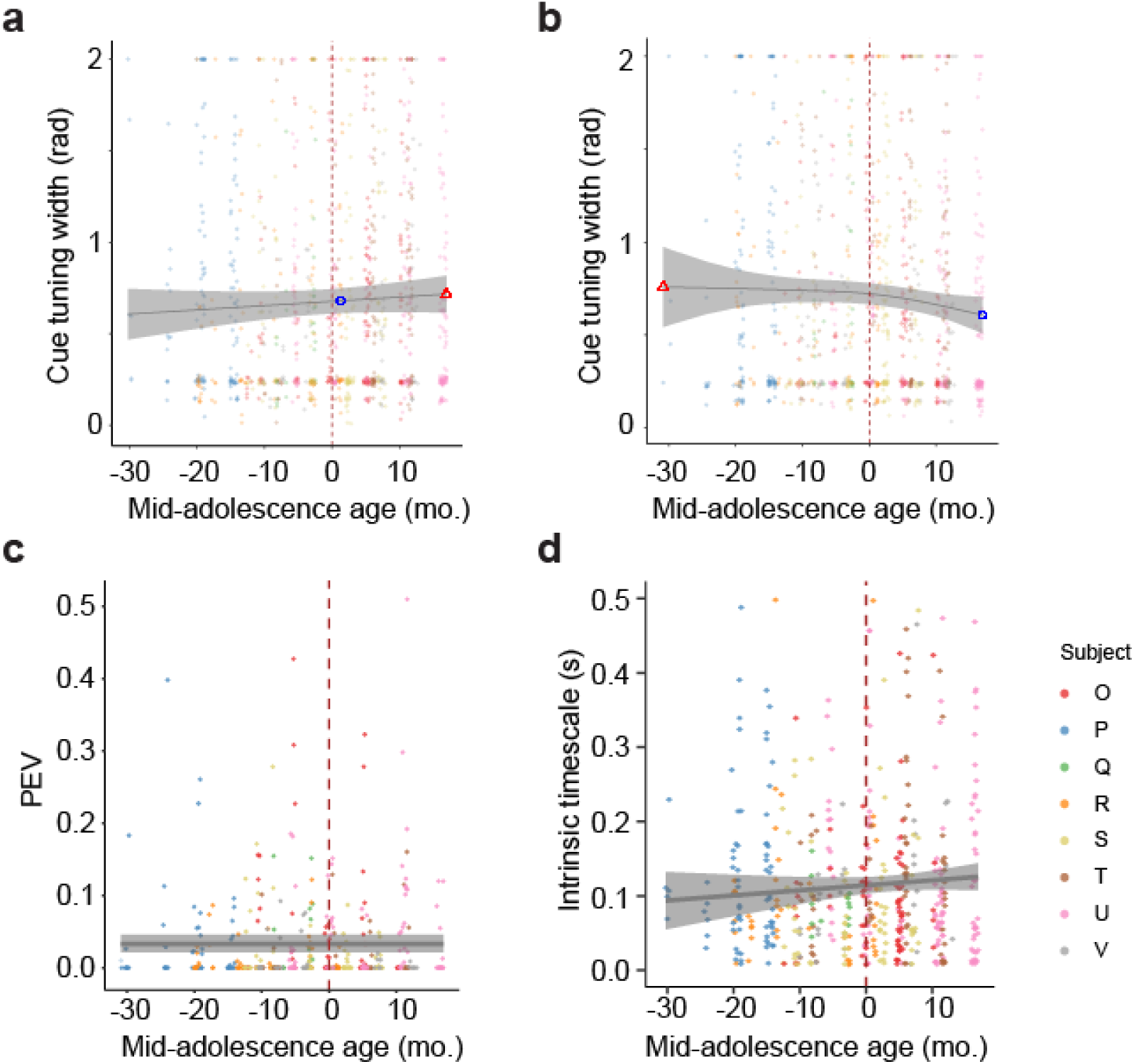
Maturation of neuronal tuning and intrinsic timescale. **a.** Tuning width estimated during the cue period as a function of time relative to the mid-adolescence age. Gray line showed the GAMM fitted trajectory. Gray shaded regions denote the 95% confidence intervals (CIs). The dashed vertical line denotes the mid-adolescence age 0. **b.** As in a, for tuning during the delay period of the ODR task. **c.** Percentage of explained variance (ω^2^) of each neuron during cue epoch of ODR task. **d.** Intrinsic timescale as a function of time relative to the mid-adolescence age, each dot is one neuron (n = 508).

**Extended Data Fig. 4.**
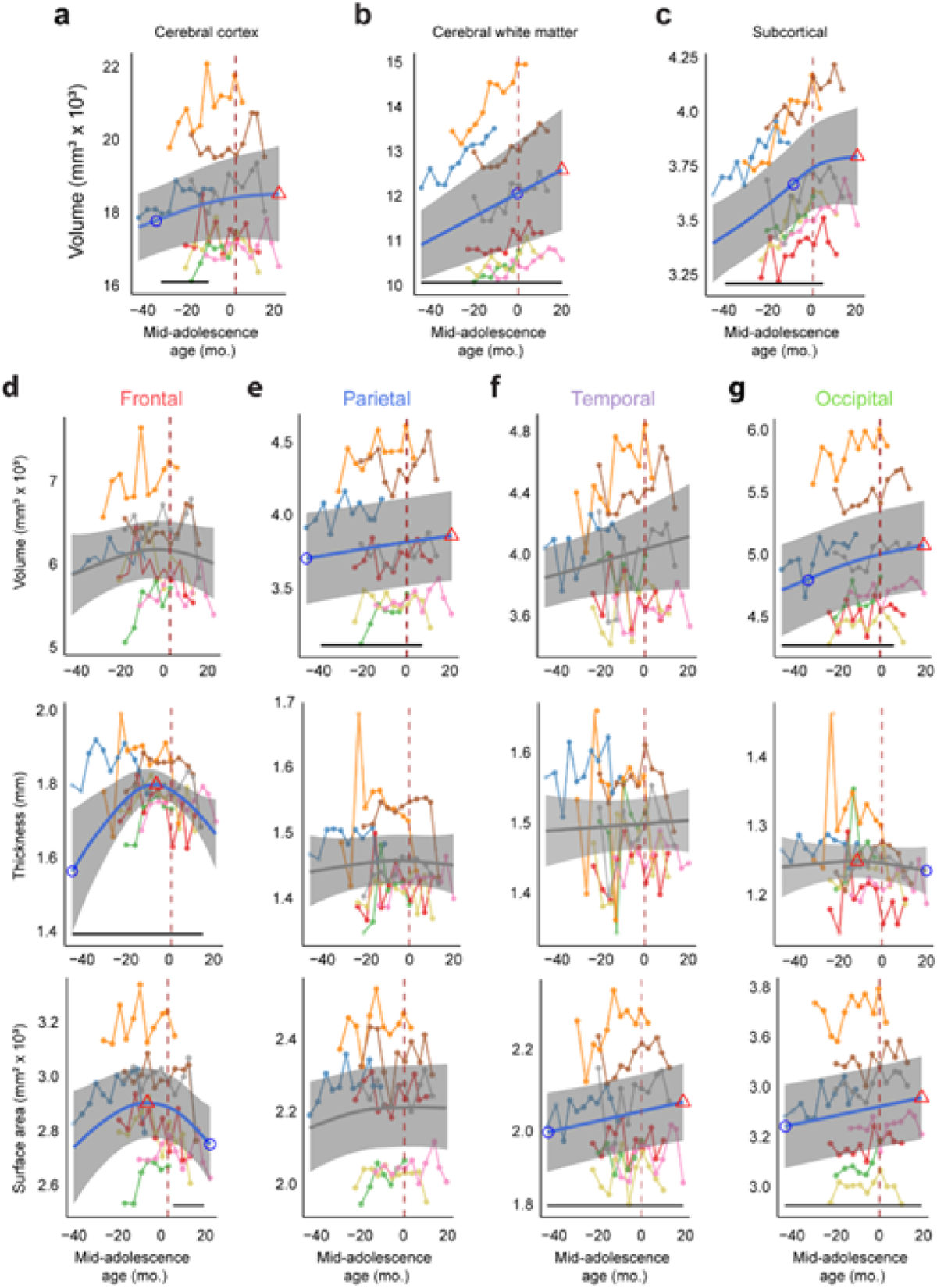
Raw data and fitted structural developmental trajectories of brain segmentations and lobes. (a) Total gray matter (cerebral cortical) volume as a function of age. Blue or gray curve indicates the GAMM fitted trajectory. Gray shaded regions denote the 95% confidence intervals (CIs). Blue circle denotes the time of peak development velocity. Red triangle denotes the time of maximum value. Dashed vertical line denotes the mid-adolescence age 0. The horizontal bar denotes significant developmental effect intervals. (b) As in a, for white matter volume as a function of age. (c) As in a, for subcortical volume as a function of age. (d-g)Volume, thickness and surface area of the four lobes of the cerebral cortex as a function of age. In each panel, top: cortical volume; middle: cortical thickness; bottom: surface area. ROIs were determined using Charm atlas level 1.

**Extended Data Fig. 5.**
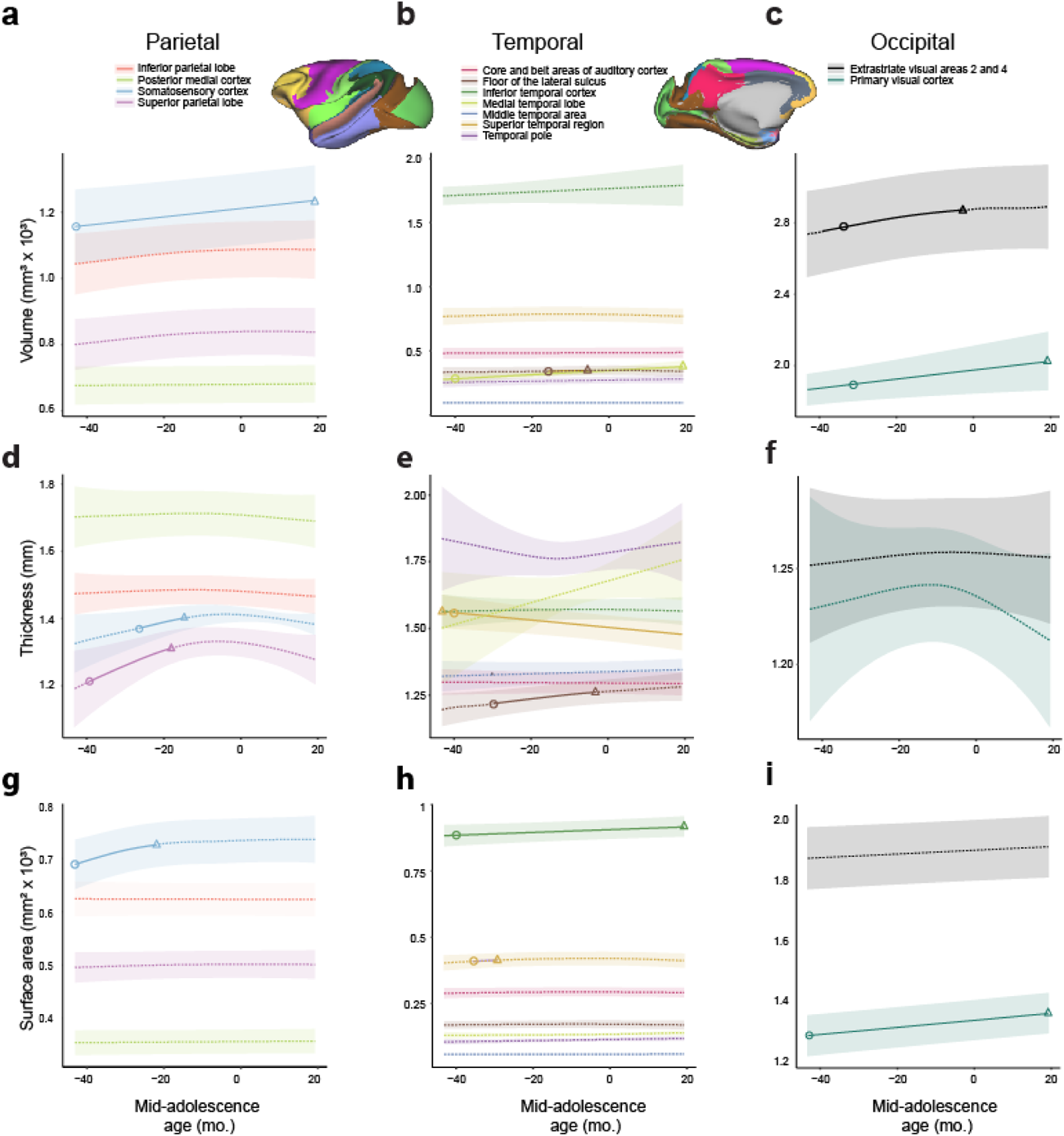
Structural changes of areas during development in parietal, temporal and occipital lobes. (a, d, g) (a) cortical volume, (d) cortical thickness and (g) surface area of areas in parietal lobe as a function of age. (b, e, h) (b) cortical volume, (e) cortical thickness and (h) surface area of areas in temporal lobe as a function of age. (c, f, i) (c) cortical volume, (f) cortical thickness and (i) surface area of areas in occipital lobe as a function of age. Each curve indicates the GAMM fitted trajectory of one area in Charm 2 atlas in each lobe. Solid curves indicate significant developmental effect of the GAMM fittings. Shaded ribbons denote the 95% confidence intervals (CIs). Circle denotes the time of peak development velocity. Triangle denotes the time of maximum volume.

**Extended Data Fig. 6.**
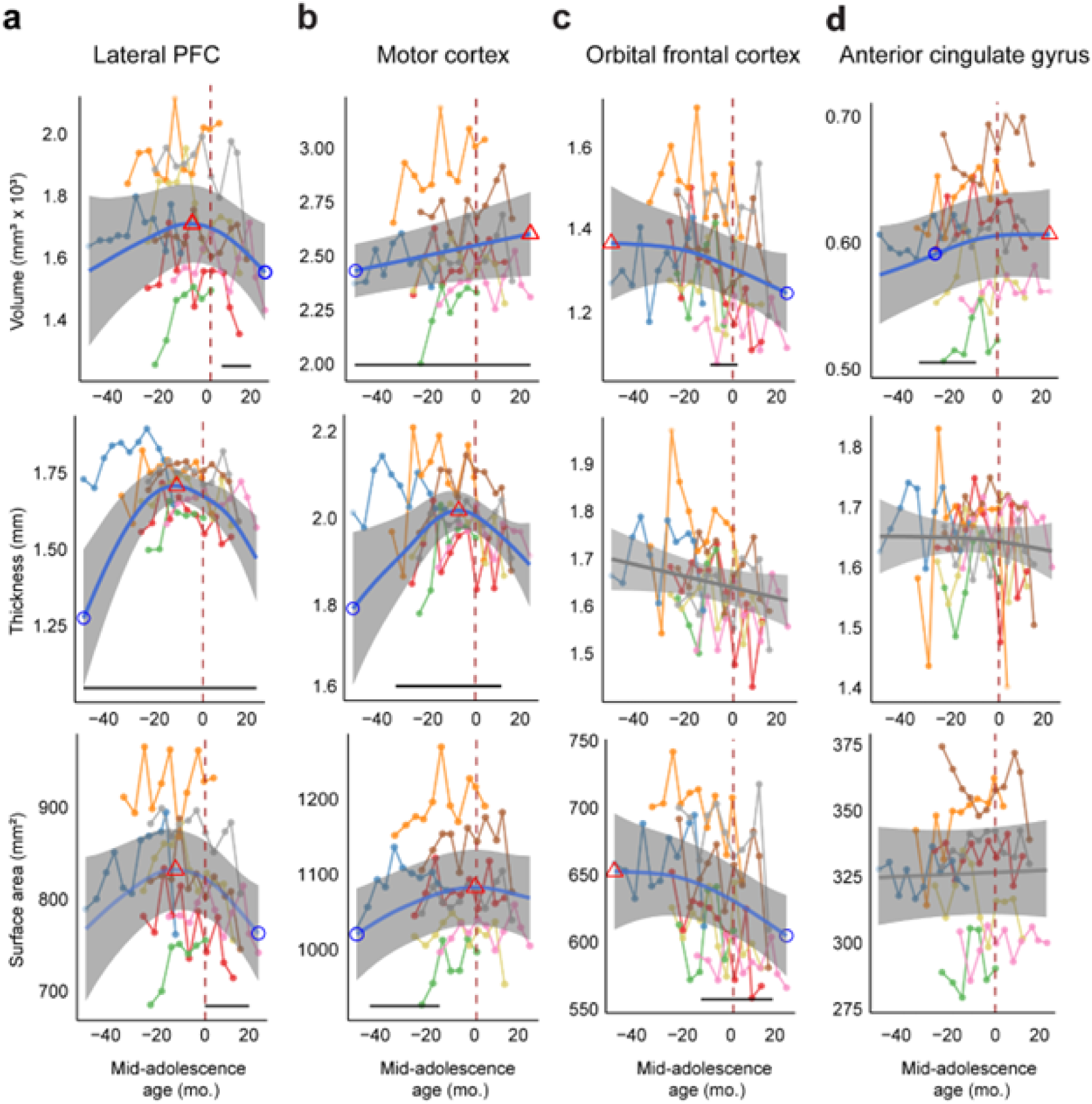
Raw data and fitted structural developmental trajectories of areas in frontal lobe. (a-d) Volume, thickness and surface area of the areas in frontal lobe as a function of age. In each panel, top: cortical volume; middle: cortical thickness; bottom: surface area. ROIs were determined using Charm atlas level 2. Blue or gray curve indicates the GAMM fitted trajectory. Gray shaded regions denote the 95% confidence intervals (CIs). Blue circle denotes the time of peak development velocity. Red triangle denotes the time of maximum value. Dashed vertical line denotes the mid-adolescence age 0. The horizontal bar denotes significant developmental effect intervals.

**Extended Data Fig. 7.**
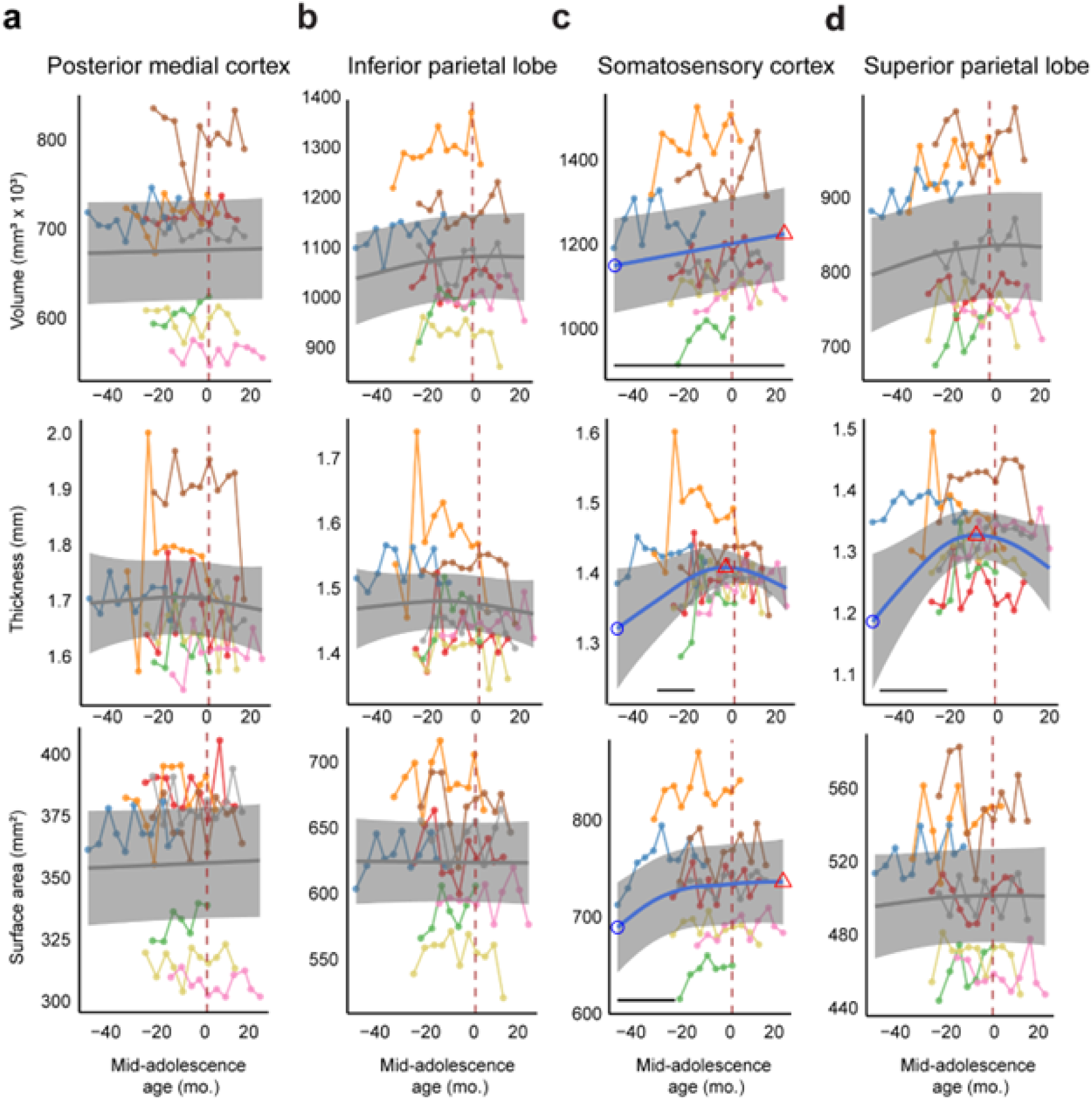
Raw data and fitted structural developmental trajectories of areas in parietal lobe. (a-d) Volume, thickness and surface area of the areas in parietal lobe as a function of age. In each panel, top: cortical volume; middle: cortical thickness; bottom: surface area. ROIs were determined using Charm atlas level 2. Blue or gray curve indicates the GAMM fitted trajectory. Gray shaded regions denote the 95% confidence intervals (CIs). Blue circle denotes the time of peak development velocity. Red triangle denotes the time of maximum value. Dashed vertical line denotes the mid-adolescence age 0. The horizontal bar denotes significant developmental effect intervals.

**Extended Data Fig. 8.**
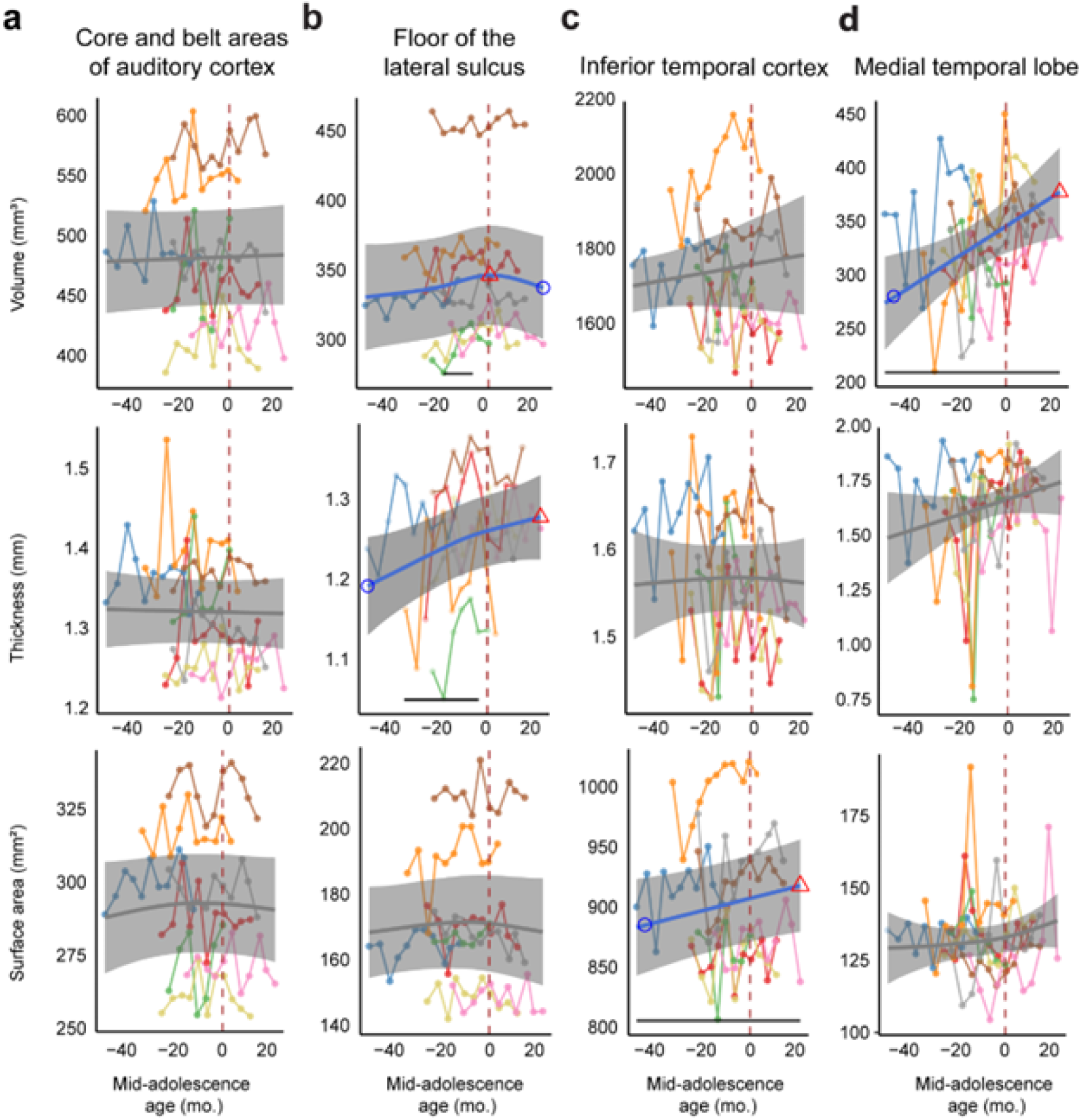
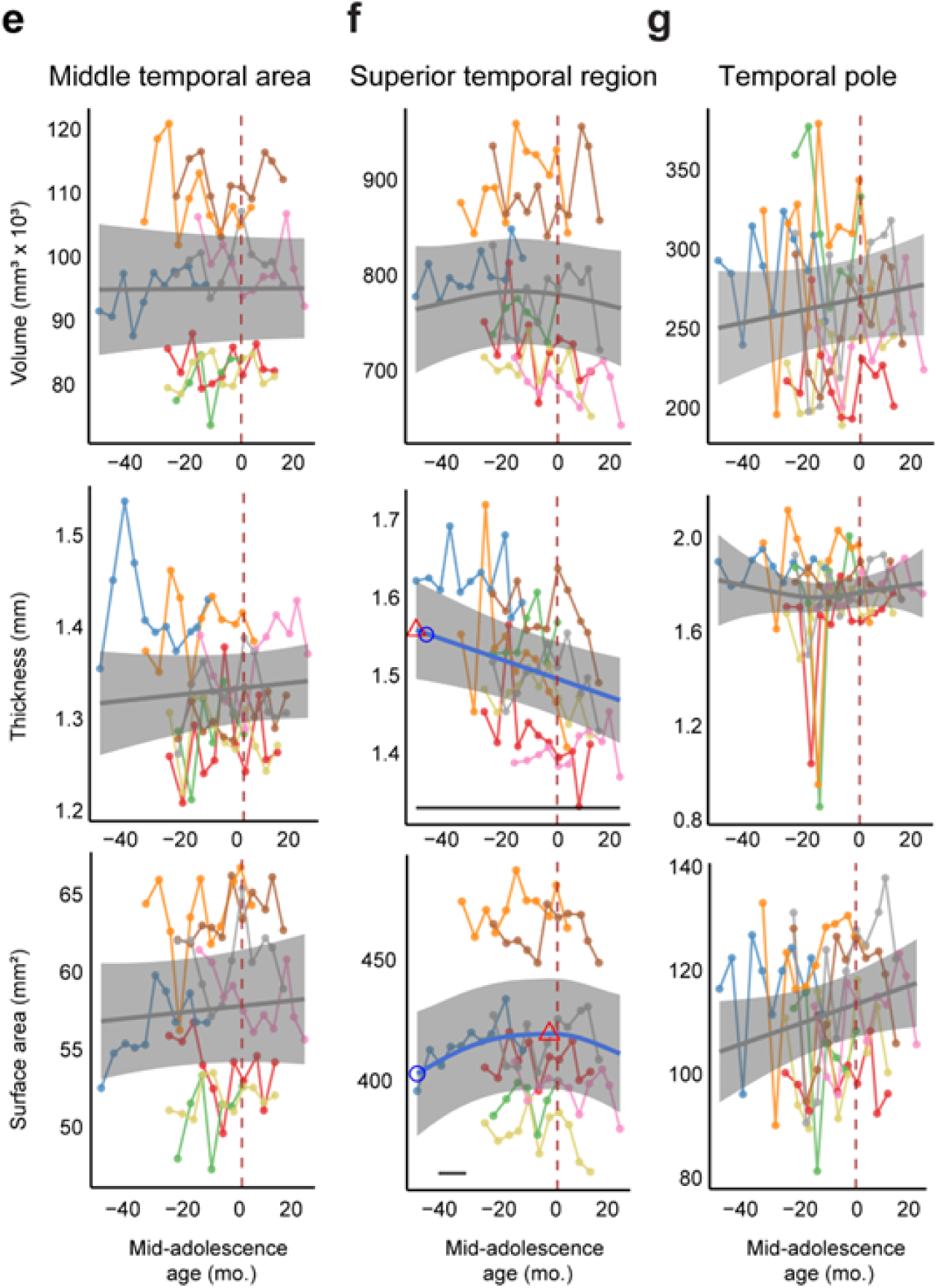
Raw data and fitted structural developmental trajectories of areas in temporal lobe. (a-g) Volume, thickness and surface area of the areas in temporal lobe as a function of age. In each panel, top: cortical volume; middle: cortical thickness; bottom: surface area. ROIs were determined using Charm atlas level 2. Blue or gray curve indicates the GAMM fitted trajectory. Gray shaded regions denote the 95% confidence intervals (CIs). Blue circle denotes the time of peak development velocity. Red triangle denotes the time of maximum value. Dashed vertical line denotes the mid-adolescence age 0. The horizontal bar denotes significant developmental effect intervals.

**Extended Data Fig. 9.**
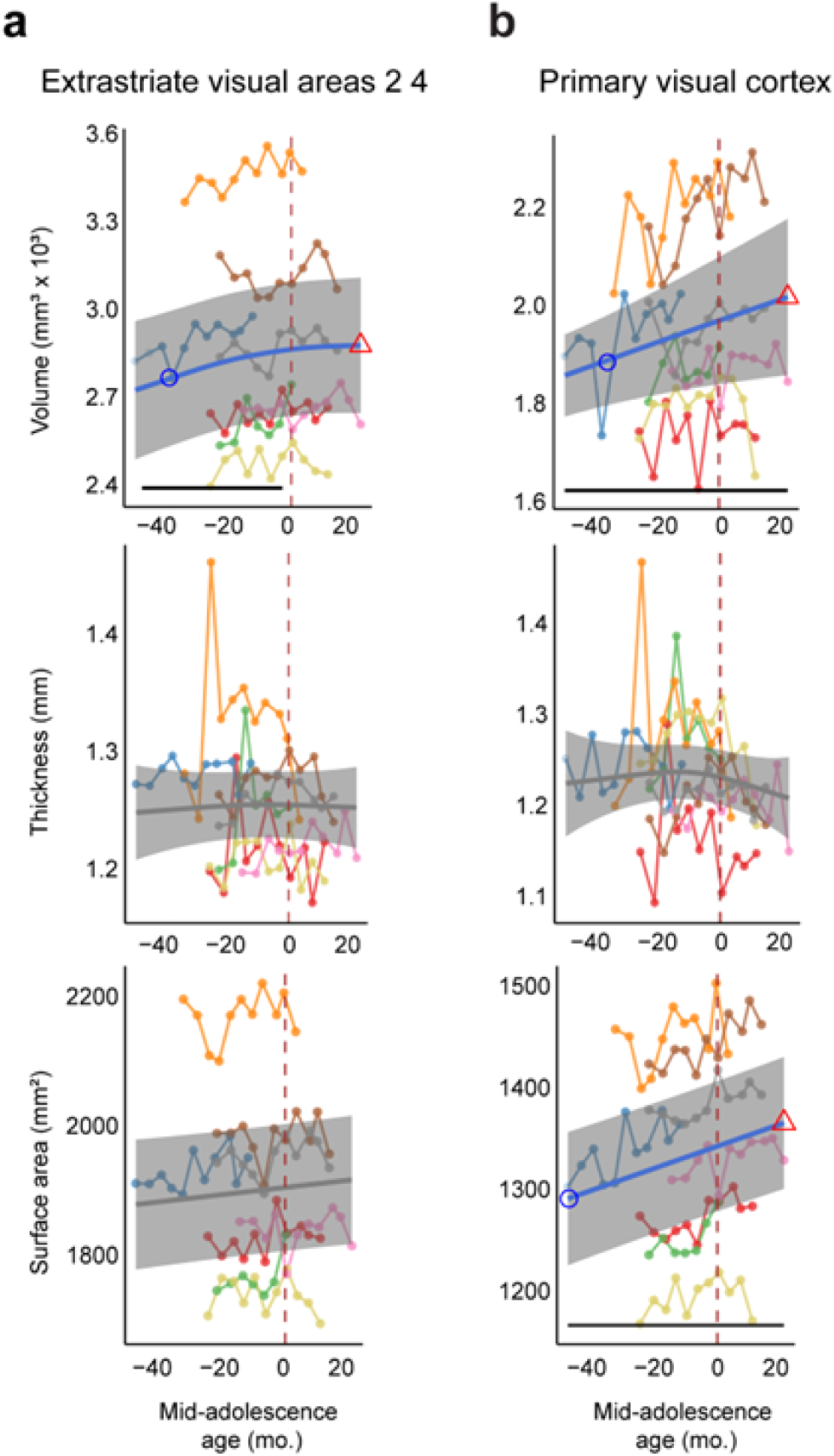
Raw data and fitted structural developmental trajectories of areas in occipital lobe. (a-d) Volume, thickness and surface area of the areas in occipital lobe as a function of age. In each panel, top: cortical volume; middle: cortical thickness; bottom: surface area. ROIs were determined using Charm atlas level 2. Blue or gray curve indicates the GAMM fitted trajectory. Gray shaded regions denote the 95% confidence intervals (CIs). Blue circle denotes the time of peak development velocity. Red triangle denotes the time of maximum value. Dashed vertical line denotes the mid-adolescence age 0. The horizontal bar denotes significant developmental effect intervals.

**Extended Data Fig. 10.**
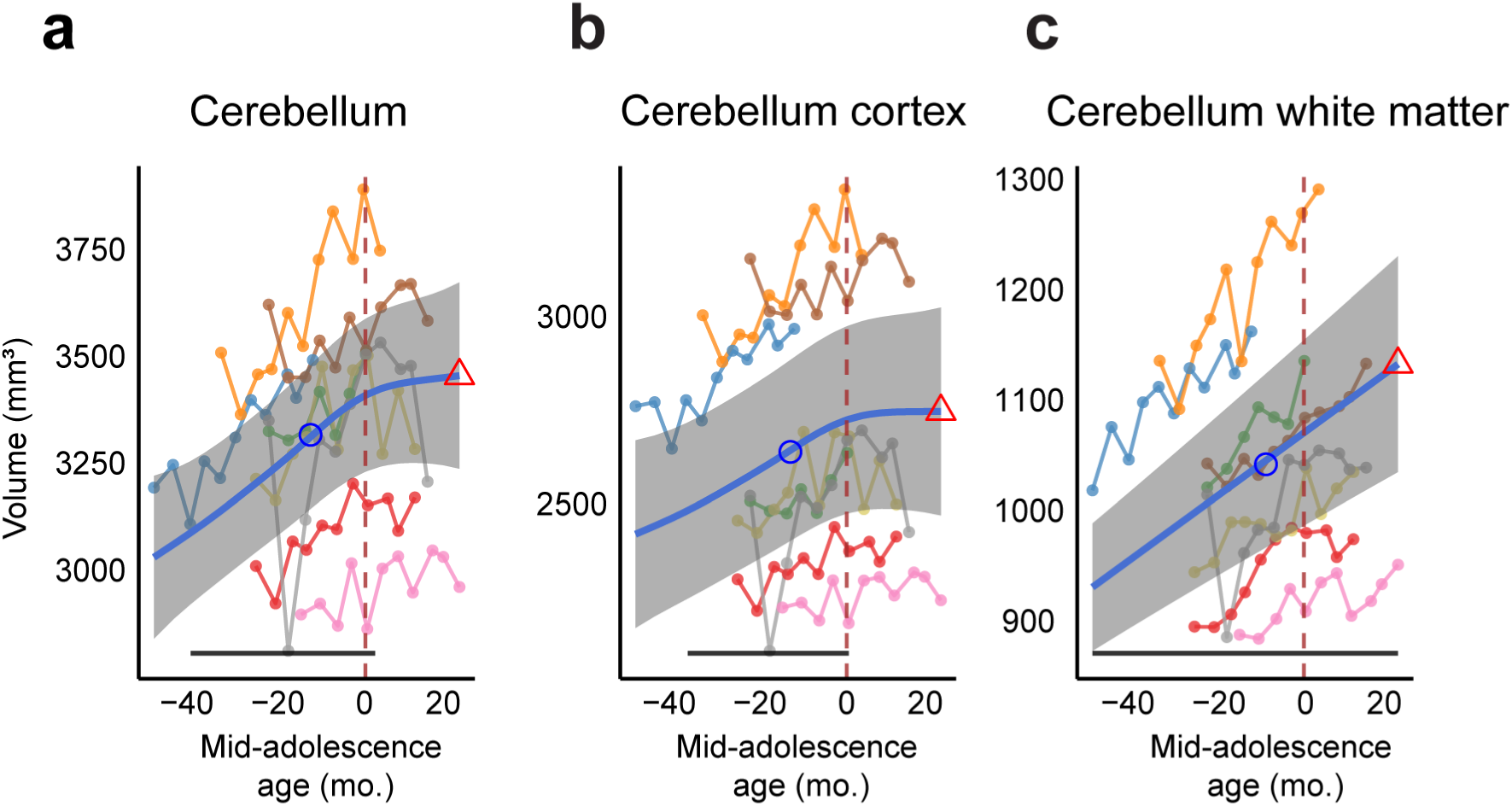
Raw data and fitted volumetric developmental trajectories of cerebellum. (a-c) Volume of cerebellum, cerebellum cortex and cerebellum white matter as a function of age. In each panel. Blue curve indicates the GAMM fitted trajectory. Gray shaded regions denote the 95% confidence intervals (CIs). Blue circle denotes the time of peak development velocity. Red triangle denotes the time of maximum value. Dashed vertical line denotes the mid-adolescence age 0. The horizontal bar denotes significant developmental effect intervals.

**Extended Data Fig. 11.**
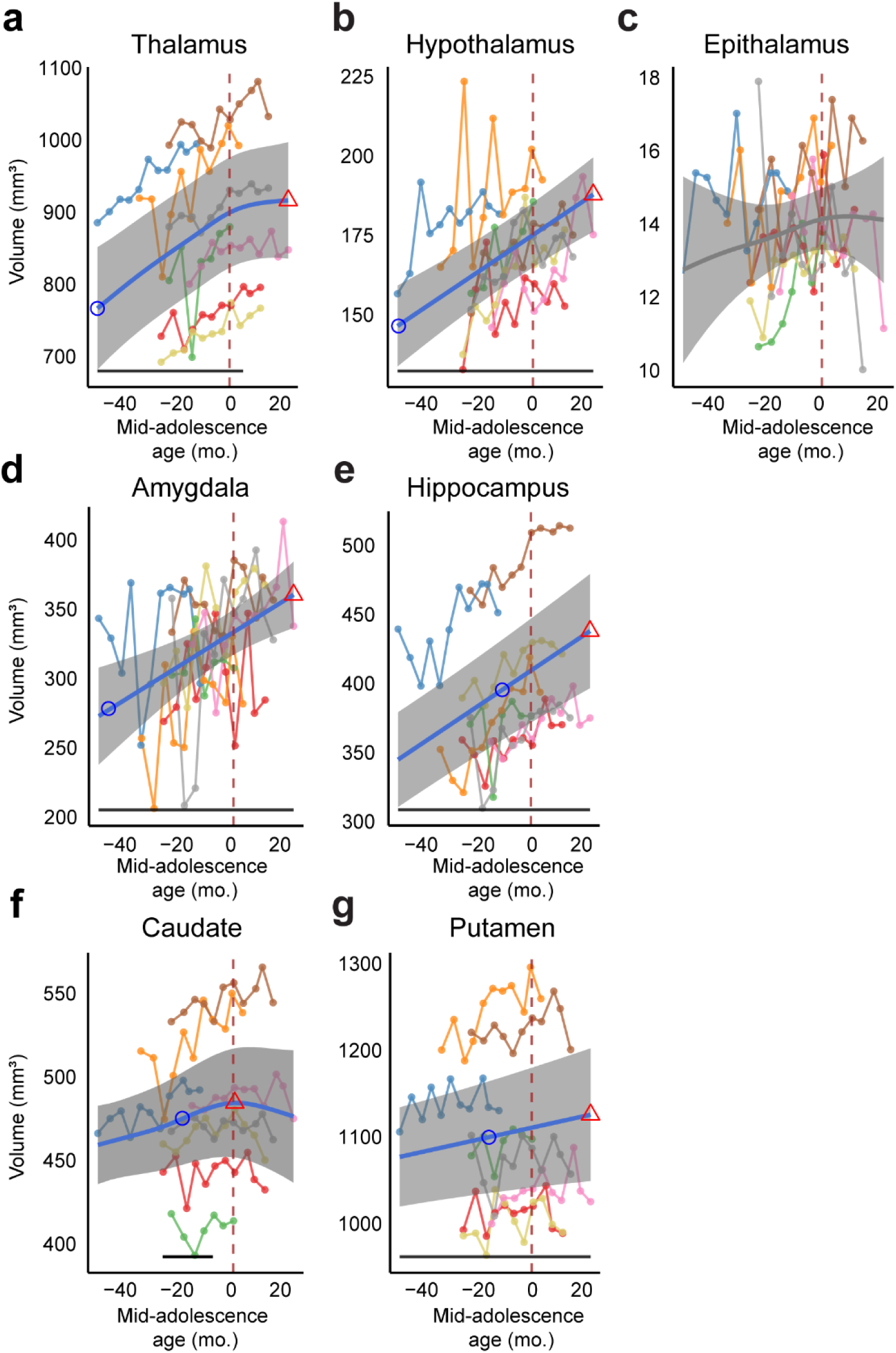
Raw data and fitted volumetric developmental trajectories of subcortical regions. Blue curve indicates the GAMM fitted trajectory. Gray shaded regions denote the 95% confidence intervals (CIs). Blue circle denotes the time of peak development velocity. Red triangle denotes the time of maximum value. Dashed vertical line denotes the mid-adolescence age 0. The horizontal bar denotes significant developmental effect intervals.

**Extended Data Fig. 12.**
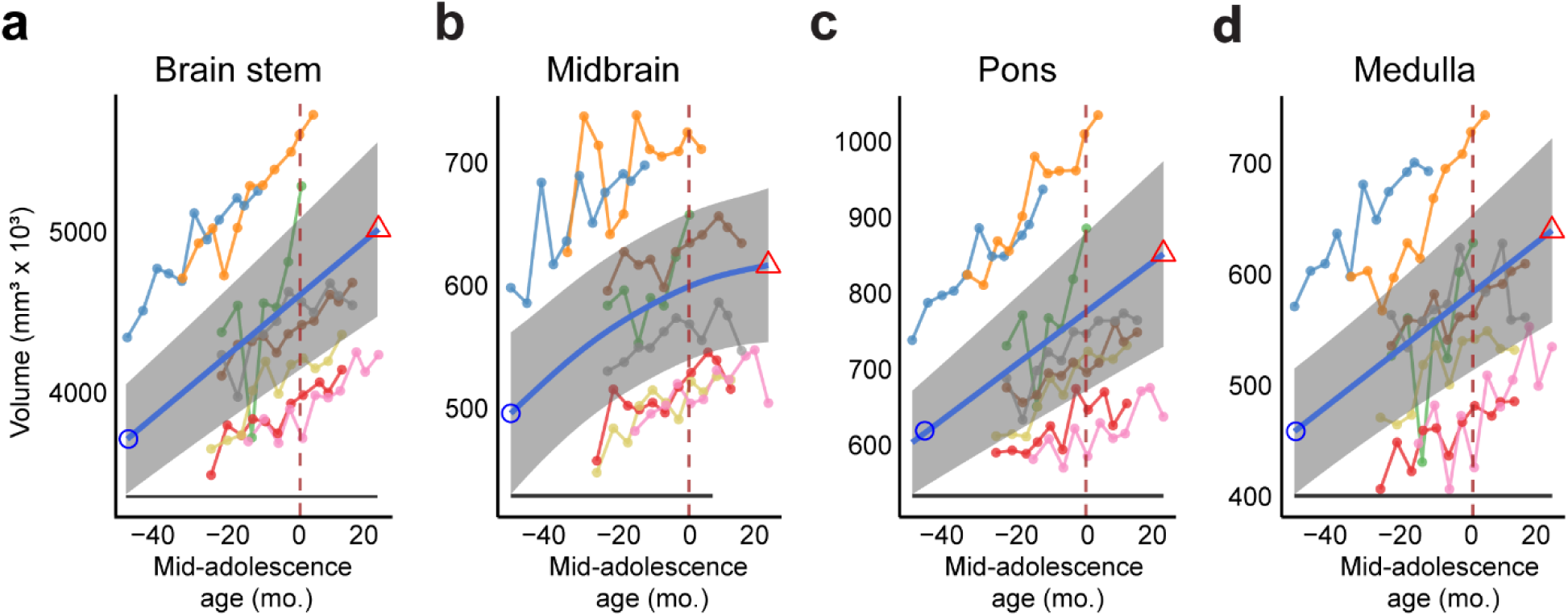
Raw data and fitted volumetric developmental trajectories of Brain stem structures. Blue curve indicates the GAMM fitted trajectory. Gray shaded regions denote the 95% confidence intervals (CIs). Blue circle denotes the time of peak development velocity. Red triangle denotes the time of maximum value. Dashed vertical line denotes the mid-adolescence age 0. The horizontal bar denotes significant developmental effect intervals.

**Extended Data Fig. 13.**
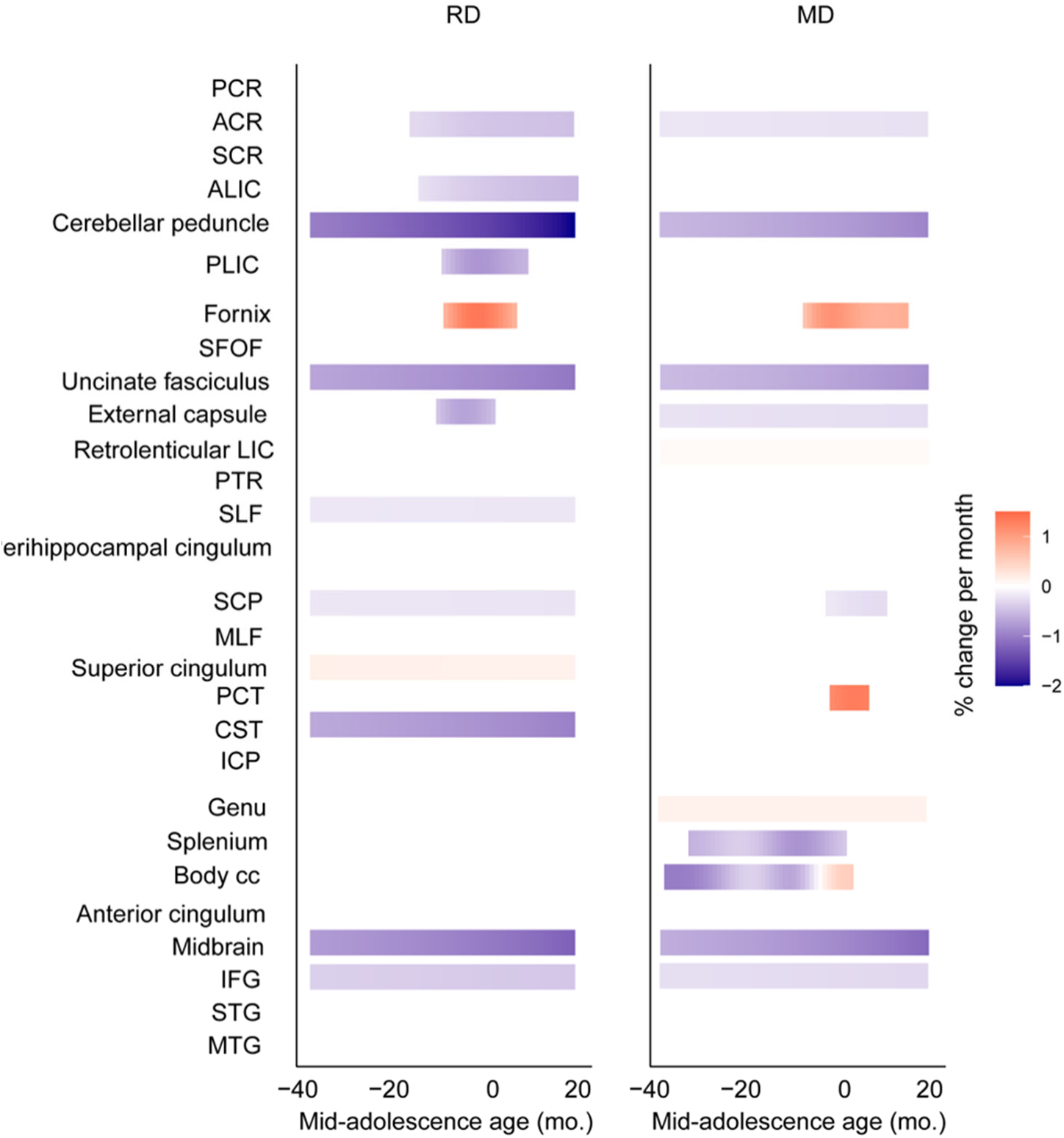
White matter maturation during development. Similar to Figure 5g, Stages of significant growth and timing of maturation of Radial Diffusivity (RD) and Mean Diffusivity (MD) of major white matter tracts. Each row is an ROI grouped to projection, association and commissural tracts, and brainstem white matter regions and short-range white matter. Rows are sorted in the same order as in Figure 5g, according to their time of maturation of FA in each group. Colors represent % change per month (red = increase, blue = decrease).

**Extended Data Fig. 14.**
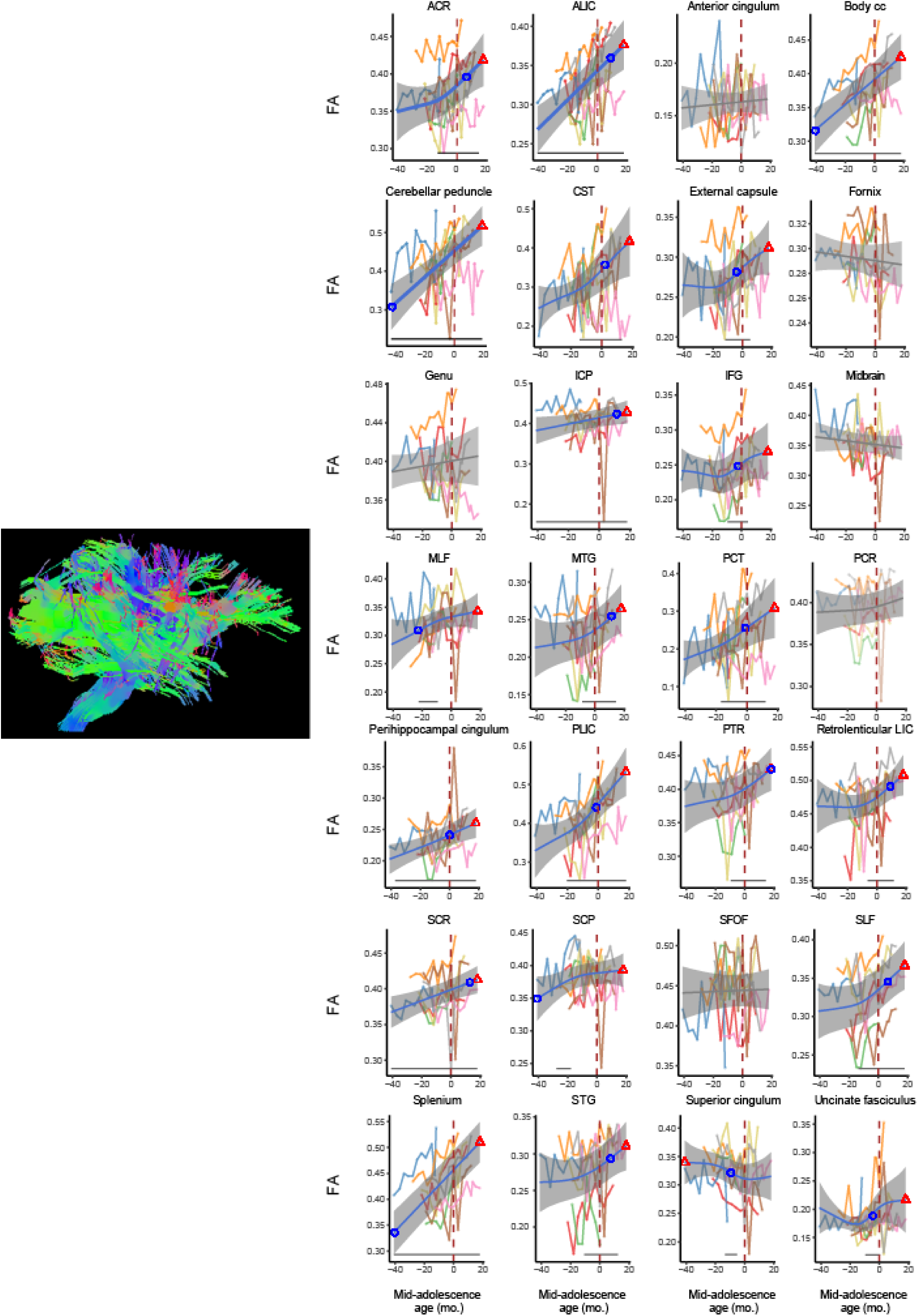
Raw data and fitted FA developmental trajectories. Left, atlas of tracts analyzed. Whole-brain tractography pathways were color-coded where red represents transverse (left to right) fibers, blue for craniocaudal (dorsal to ventral) fibers, and green for anteroposterior (anterior to posterior) fibers. Right, developmental trajectories. Blue or gray curve indicates the GAMM fitted trajectory. Gray shaded regions denote the 95% confidence intervals (CIs). Blue circle denotes the time of peak development velocity. Red triangle denotes the time of maximum value. Dashed vertical line denotes the mid-adolescence age 0. The horizontal bar denotes significant developmental effect intervals. Panels were sorted in alphabetical order.

**Extended Data Fig. 15.**
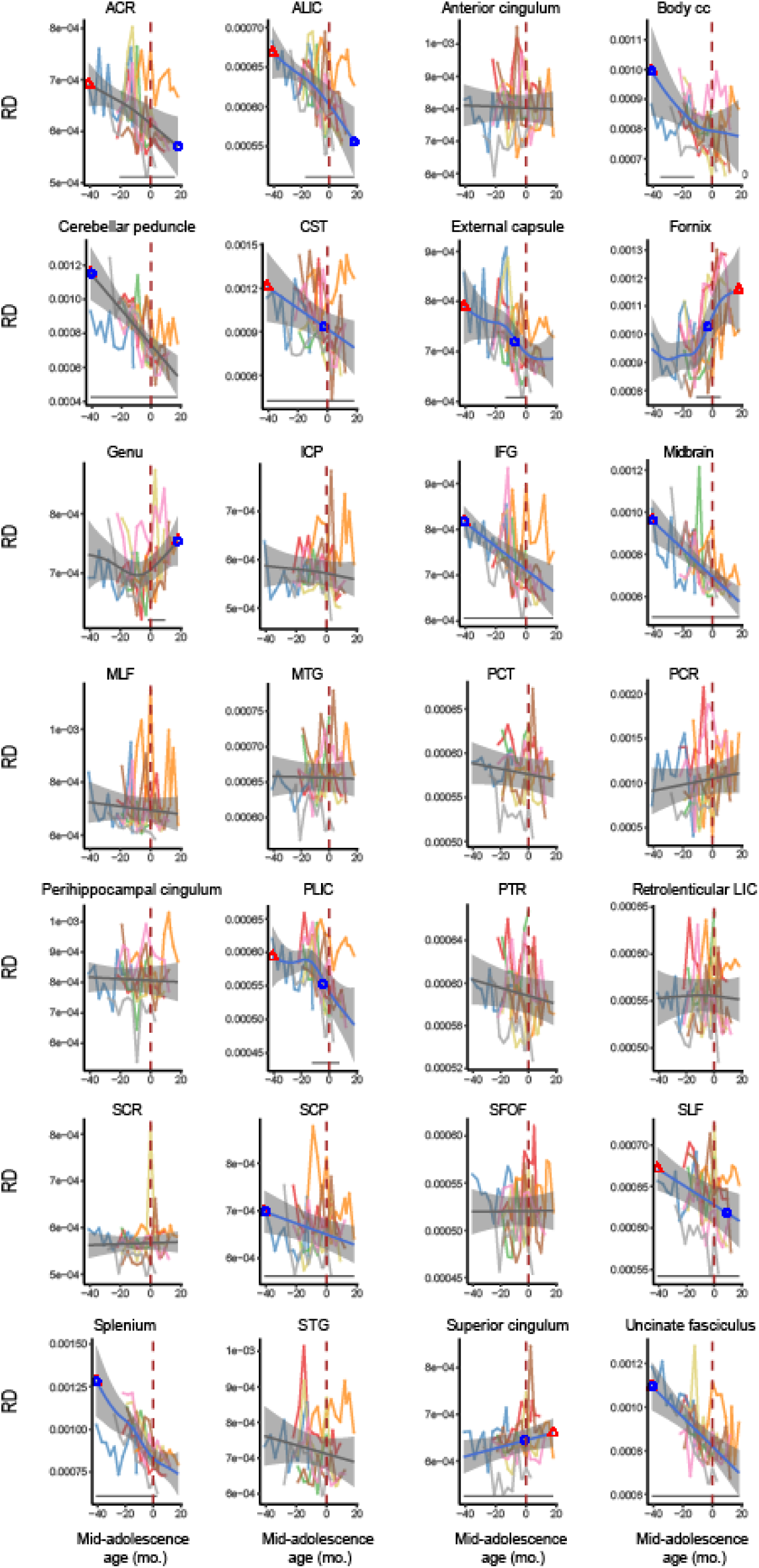
Raw data and fitted RD developmental trajectories. Blue or gray curve indicates the GAMM fitted trajectory. Gray shaded regions denote the 95% confidence intervals (CIs). Blue circle denotes the time of peak development velocity. Red triangle denotes the time of maximum value. Dashed vertical line denotes the mid-adolescence age 0. The horizontal bar denotes significant developmental effect intervals. Panels were sorted in alphabetical order.

**Extended Data Fig. 16.**
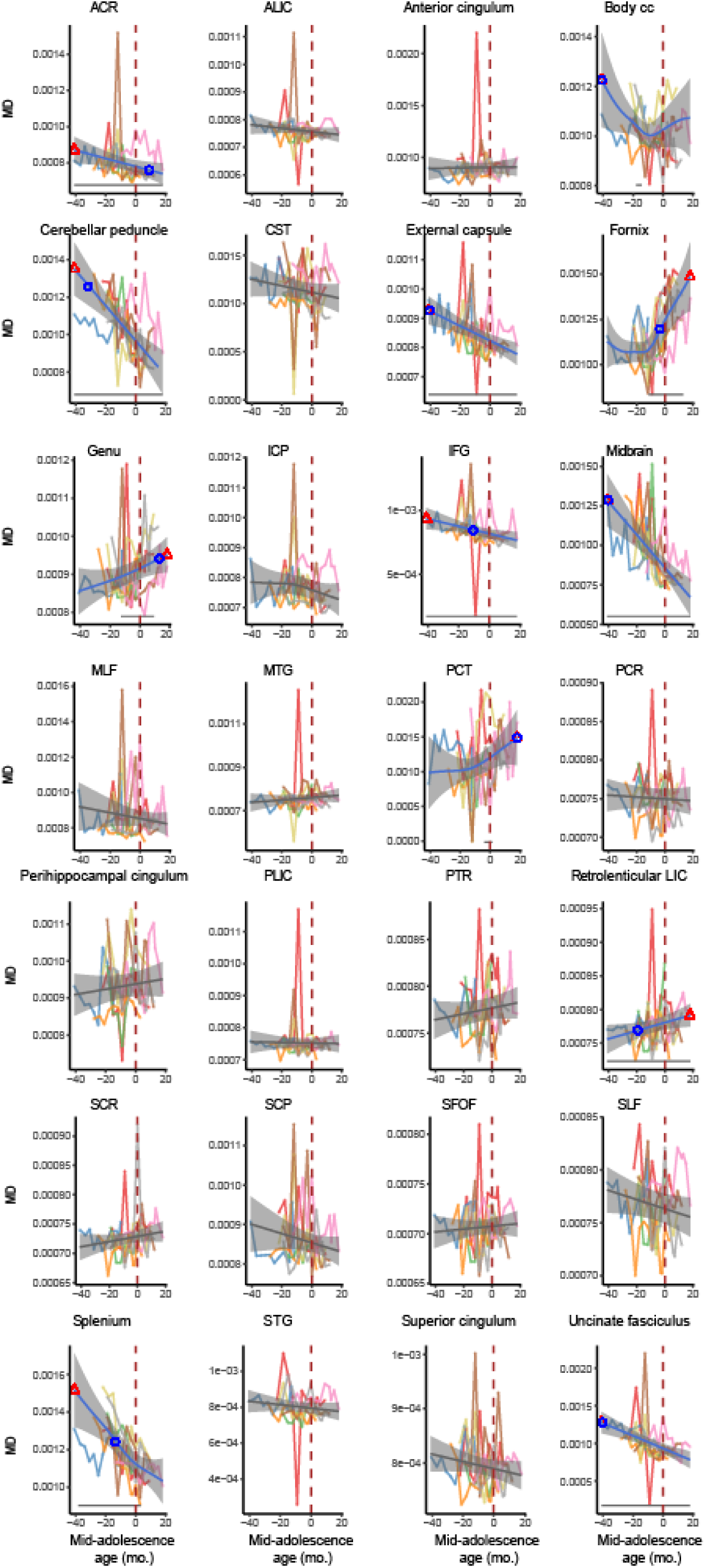
Raw data and fitted MD developmental trajectories. Blue or gray curve indicates the GAMM fitted trajectory. Gray shaded regions denote the 95% confidence intervals (CIs). Blue circle denotes the time of peak development velocity. Red triangle denotes the time of maximum value. Dashed vertical line denotes the mid-adolescence age 0. The horizontal bar denotes significant developmental effect intervals. Panels were sorted in alphabetical order.

**Extended Data Fig. 17.**
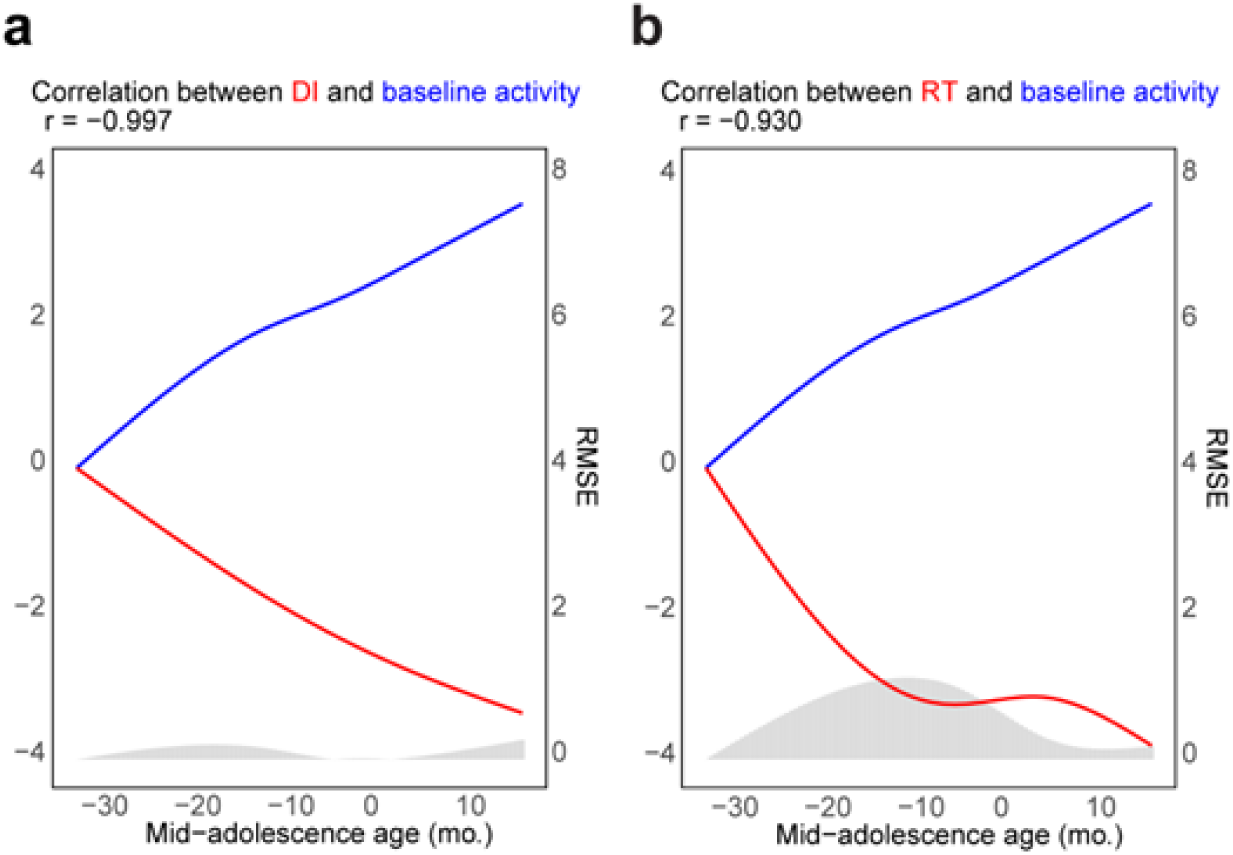
Correlation between the GAMM fitted trajectories of behavioral measures and baseline activity of PFC neurons. (a) Correlation between the GAMM fitted trajectories of DI and baseline activity of PFC neurons. Curves represent normalized GAMM fitted trajectories. Histogram indicates RMSE between corresponding prediction values of the GAMMs. (b) As in (a), for the GAMM fitted trajectories of RT and baseline activity of PFC neurons.

**Extended Data Fig. 18.**
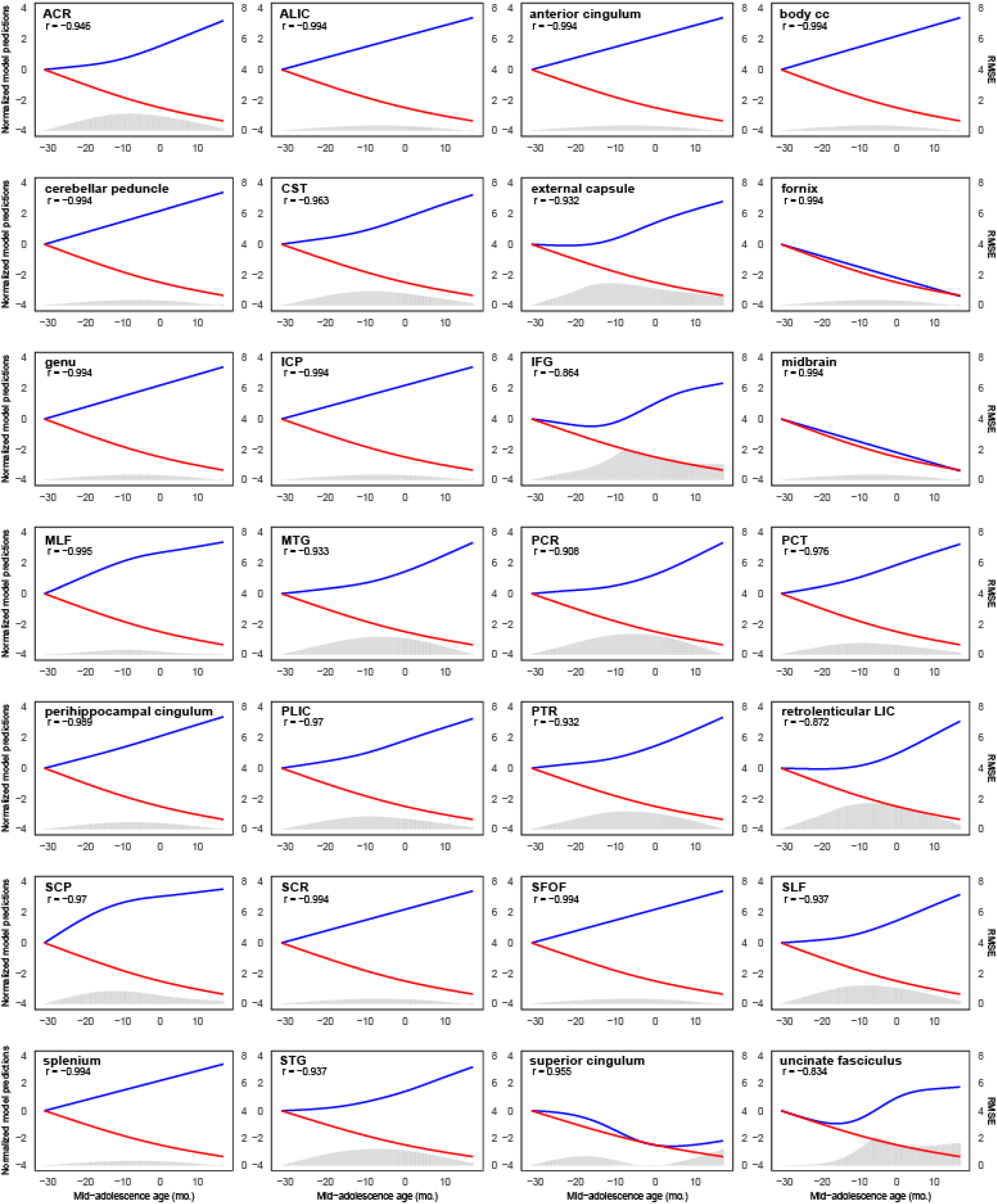
Correlation between the GAMM fitted trajectories of DI and FA of each white matter tracts. In each panel, blue curve represents normalized GAMM fitted trajectory of the FA of the tract and red curve represents normalized GAMM fitted trajectory of DI in ORD task. Histogram indicates RMSE between corresponding prediction values of the GAMMs.

**Extended Data Fig. 19.**
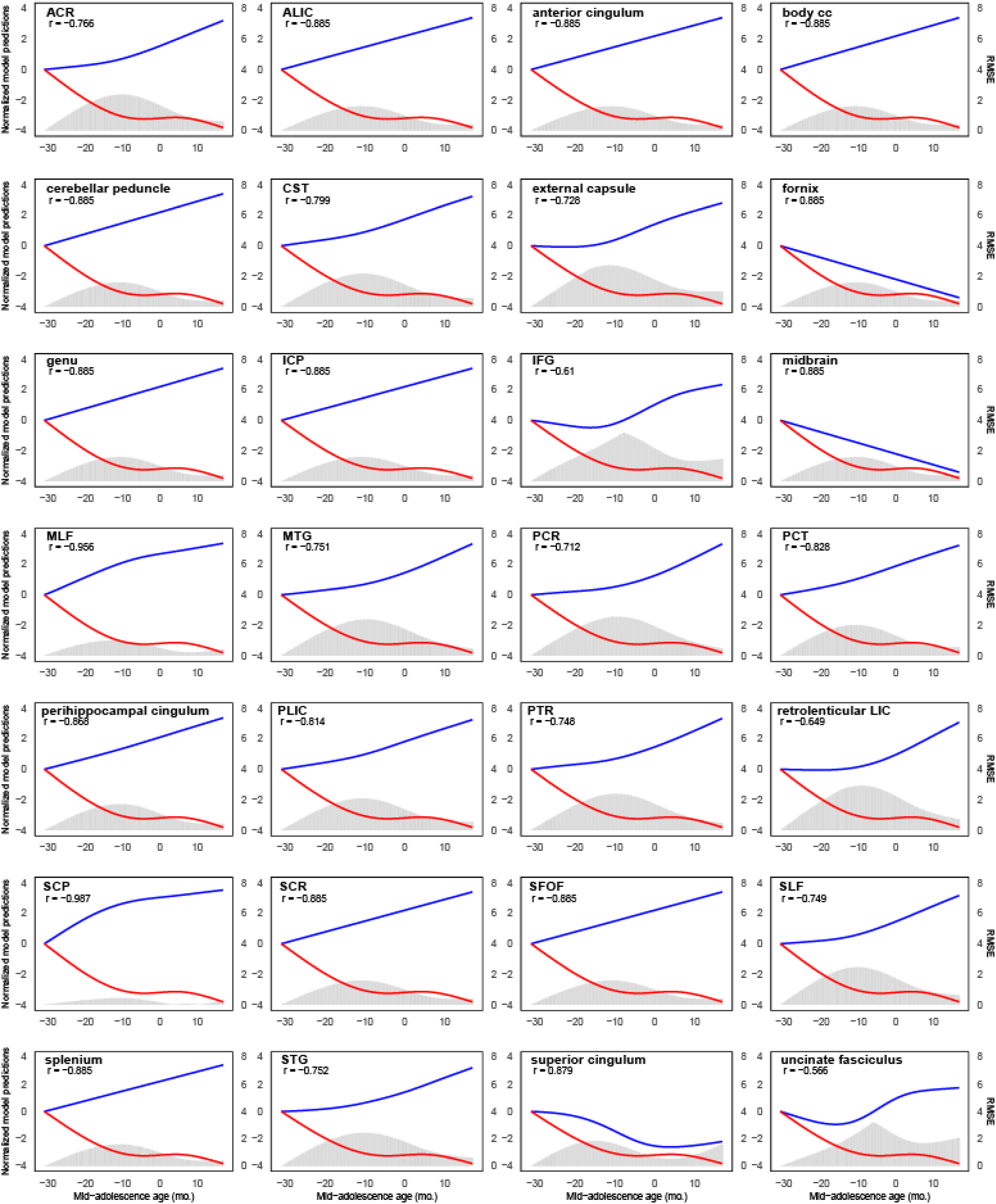
Correlation between the GAMM fitted trajectories of RT and FA of each white matter tracts. In each panel, blue curve represents normalized GAMM fitted trajectory of the FA of the tract and red curve represents normalized GAMM fitted trajectory of RT in ORD task. Histogram indicates RMSE between corresponding prediction values of the GAMMs.

**Extended Data Table 1.** Morphometric measures and model fit parameters aligned on chronological age and mid-adolescence age.

| Measures | Variable | edf | F | P value | AIC |
| --- | --- | --- | --- | --- | --- |
| Body weight | Chronological age | 3.14 | 22.14 | $<10^{-6}$ | 834.10 |
| | **mid-adolescence age** | 5.67 | 26.77 | $<10^{-6}$ | **792.43** |
| Canine length | Chronological age | 1.00 | 153.85 | $<10^{-6}$ | 1054.95 |
| | **mid-adolescence age** | 4.51 | 39.20 | $<10^{-6}$ | **1035.80** |
| Femur length | **Chronological age** | 6.27 | 29.68 | $<10^{-6}$ | **753.19** |
| | mid-adolescence age | 5.72 | 13.78 | $<10^{-6}$ | 771.09 |
| Nipple length | Chronological age | 4.16 | 12.88 | $4.00 \times 10^{-6}$ | 260.70 |
| | **mid-adolescence age** | 4.03 | 11.05 | $1.51 \times 10^{-5}$ | **260.32** |
| Testis length | Chronological age | 3.55 | 26.73 | $<10^{-6}$ | 677.16 |
| | **mid-adolescence age** | 5.89 | 63.84 | $<10^{-6}$ | **648.43** |
| Testis volume | Chronological age | 4.04 | 88.10 | $<10^{-6}$ | 808.09 |
| | **mid-adolescence age** | 5.40 | 97.21 | $<10^{-6}$ | **780.90** |
| Trunk length | Chronological age | 5.16 | 23.23 | $<10^{-6}$ | 692.97 |
| | **mid-adolescence age** | 5.84 | 34.28 | $<10^{-6}$ | **688.83** |

**Extended Data Table 2.** Sample sizes for neurons and sessions in each subject.

| Subject | N(neuron) | N(session) |
| --- | --- | --- |
| O | 292 | 53 |
| P | 264 | 52 |
| Q | 62 | 16 |
| R | 236 | 56 |
| S | 330 | 61 |
| T | 262 | 36 |
| U | 527 | 75 |
| V | 158 | 38 |

**Extended Data Table 3.**
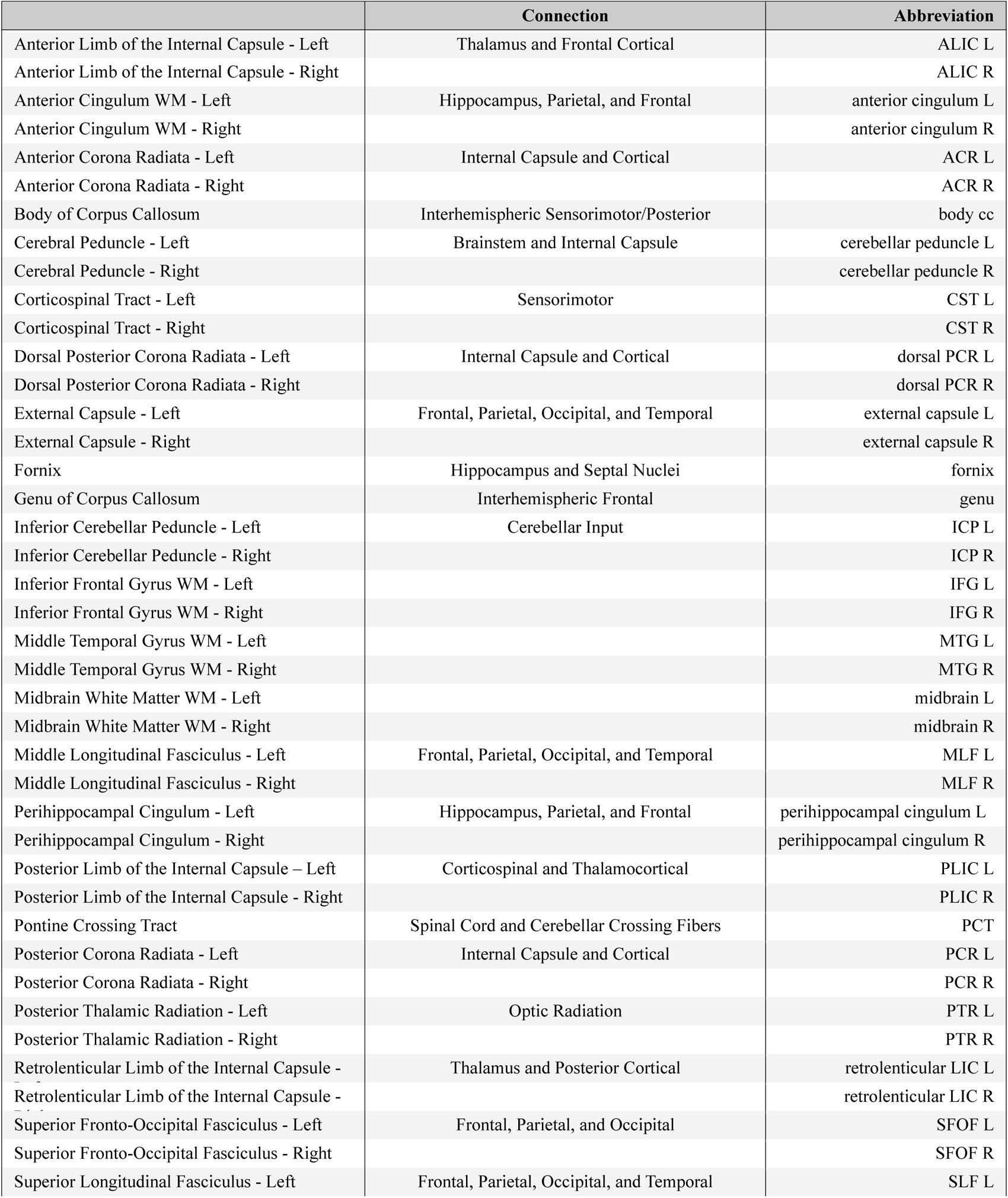
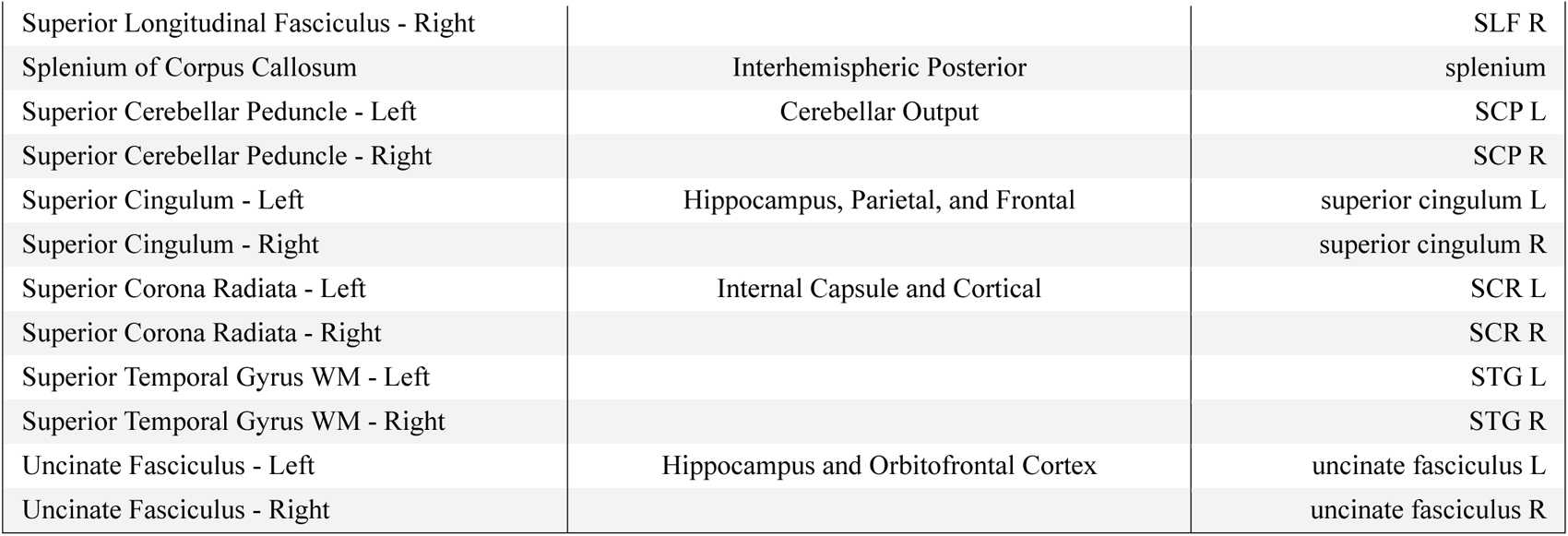
Description of selected ROIs from white matter (WM) atlas analyzed.

